# Non-canonical NF-κB signaling promotes intestinal inflammation by restraining the tolerogenic β-catenin-Raldh2 axis in dendritic cells

**DOI:** 10.1101/2023.12.03.569755

**Authors:** Alvina Deka, Naveen Kumar, Meenakshi Chawla, Namrata Bhattacharya, Sk Asif Ali, Swapnava Basu, Bhawna, Upasna Madan, Shakti Kumar, Bhabatosh Das, Debarka Sengupta, Amit Awasthi, Soumen Basak

**Author notes:** Correspondence should be addressed to S.B. These authors have contributed equally to the work (Alvina Deka, Naveen Kumar).

## Abstract

Dendritic cell (DC) dysfunctions exacerbate intestinal pathologies. However, the mechanisms compromising DC-mediated immune controls remain unclear. We found that intestinal DCs from mice subjected to experimental colitis possessed heightened non-canonical NF-κB signaling, which activates the RelB:p52 heterodimer. Genetic inactivation of this pathway in DCs alleviated inflammation in colitogenic mice. Unexpectedly, RelB:p52 deficiency diminished the transcription of Axin1, a critical component of the β-catenin destruction complex. This reinforced β-catenin-driven expression of Raldh2, which imparts tolerogenic DC attributes by promoting retinoic acid (RA) synthesis. Indeed, DC-specific non-canonical NF-κB impairment improved the colonic frequency of Tregs and IgA^+^ B cells, which fostered luminal IgA and eubiosis. Introducing β-catenin haploinsufficiency in non-canonical NF-κB-deficient DCs moderated Raldh2 activity, reinstating colitogenic sensitivity in mice. Finally, IBD patients displayed a deleterious non-canonical NF-κB signature in intestinal DCs. In sum, we establish a DC network that integrates non-canonical NF-κB signaling to subvert RA metabolic pathway in fueling intestinal inflammation.

**Significance (100):** Distorted dendritic cell (DC) functions have been implicated in aberrant intestinal inflammation; however, the underlying mechanism remains obscure. We discovered that the non-canonical NF-κB pathway exacerbates inflammation in the colitogenic gut by downmodulating β-catenin-driven synthesis of Raldh2 in DCs. Raldh2 represents a key enzyme involved in the production of tolerogenic retinoic acid in intestinal DCs. Beyond regulating immune genes, therefore, non-canonical NF-κB signaling appears to instruct retinoic acid-mediated control of gut health. While we illustrate a DC network integrating immune signaling and micronutrient metabolic pathways in the intestine, our finding may have broad relevance for nutritional interventions in inflammatory ailments.

**eToC:** Deka and Kumar *et al*. illustrate a DC-circuitry that exacerbates intestinal inflammation in IBD patients and colitogenic mice. Non-canonical NF-κB signaling restrains β-catenin in DCs to downmodulate Raldh2, which promotes tolerogenic RA synthesis, leading to diminished Treg and IgA^+^ cell frequencies in the gut.

**Highlights:**

- Aberrant intestinal inflammation is associated with and exacerbated by non-canonical NF-κB signaling in DCs.
- Non-canonical signaling restrains the tolerogenic β-catenin-Raldh2 axis in DCs by upregulating Axin1.
- DC-specific RelB:p52 impairment promotes β-catenin-dependent Treg accumulation in the gut.
- A DC defect of non-canonical signaling causes β-catenin-dependent increase in luminal sIgA, fostering the gut microbiome.

**One sentence:** The non-canonical NF-κB pathway fuels intestinal inflammation by waning the tolerogenic β-catenin-Raldh2-retinoic acid axis in DCs.

## Introduction

DCs orchestrate the balance between protective immune responses against pathogens and tolerance towards commensal microbes in the intestine. Upon microbial sensing, activated DCs produce a myriad of immunogenic cytokines, including IL-6, IL-12, and IL-23, which promote IFNγ-secreting Th1 and IL-17-secreting Th17 cells. These effector T cells direct inflammatory responses, containing gut infections. DCs also produce tolerogenic cytokines, such as IL-10 and TGFβ. In addition, mucosal DCs synthesize retinoic acid (RA) from dietary vitamin A or retinol^1^. These immunomodulatory factors support the generation and maintenance of inflammation-suppressive FoxP3^+^ Tregs and promote their gut homing. Furthermore, DC-derived RA and IL-10 enhance immunoglobulin class switching, accumulating IgA-secreting plasma B cells in the intestine^2,3^. While secretory IgA (sIgA) in the gut lumen protects from opportunistic gut pathogens, sIgA coating also shapes the gut microbiome in symbiosis with the host^4,5^. Thus, by calibrating mucosal responses, DCs ensure gut homeostasis.

Not surprisingly, distorted DC functions have been implicated in aberrant intestinal inflammation. Ulcerative colitis (UC), a form of IBD, was associated with a marked reduction in the CD103^+^ DC subset in the intestine^6^. Colonic DCs from IBD patients or mice subjected to experimental colitis were inept at synthesizing TGFβ and RA ^6,7^. These DCs were also less proficient in generating Tregs and instead promoted Th1 and Th17 cells. Accordingly, DC-specific ablation of RXRα, an RA-activated transcription factor, aggravated colitis in mice by depleting intestinal Tregs^8^ . Similarly, CD137 signaling restricted colitis in mice by upregulating RA synthesis in DCs that altered T cell homeostasis in the intestine^9^. Furthermore, vitamin A deficiency exacerbated colitis in mice by decreasing the colonic abundance of IgA-secreting cells^10^. Notably, IgA-coated gut microbes from healthy individuals provided protection, while those from IBD patients sensitized mice to experimental colitis^11,12^. What triggers such DC dysfunction in intestinal pathologies remains unclear.

The NF-κB system comprises two interlinked canonical and non-canonical signaling arms, which play important roles in the differentiation, survival, and functioning of DCs. In particular, the canonical NF-κB pathway mediates microbial signal-responsive nuclear translocation of RelA- and cRel-containing NF-κB transcription factors, which induce the expression of immunogenic cytokines, including IL-1β, IL-12, and IL-23, in DCs^13^. Previous studies suggested that the activation of canonical NF-κB signaling in DCs worsens experimental colitis in mice^14^. On the other hand, the non-canonical NF-κB pathway is activated by a subset of TNFR superfamily members, including lymphotoxin-β receptor (LTβR), GM-CSFR, and CD40^15^. In this pathway, NF-κB inducing kinase (NIK)-dependent phosphorylation promotes processing of the NF-κB precursor protein p100 into the mature p52 subunit, leading to nuclear accumulation of RelB:p52. In a non-redundant manner to the canonical pathway, non-canonical NF-κB signaling in DCs induces the expression of IL-23, which drives Th17 responses^16,17^. NIK function in DCs was implicated in the pathogenic Th17 response underlying epidermal damage in the mouse model of psoriasis^18^. DC-intrinsic role of NIK and RelB also restricted Treg accumulation in the central nervous system in the mouse model of experimental autoimmune encephalomyelitis (EAE)^17,19^. In the intestine, NIK-dependent IL-23 expression by DCs was important for Th17-mediated immune controls of *Citrobacter rodentium*^20^. As such, NIK in DCs promoted pIgR expression in IECs involving Th17-secreted IL-17 that maintained luminal IgA levels and a gut-protective microbiome. Interestingly, *LTBR* and *NFKB2* were linked in a genome-wide association to the susceptibility loci for IBD^21^. We asked if non-canonical RelB:p52 NF-κB signaling perturbed DC functions in the inflamed gut.

Here, we report that human IBD and experimental colitis in mice are associated with intensified non-canonical NF-κB signaling in intestinal DCs. Genetic disruption of this pathway in DCs alleviated chemical-induced colitis in mice. Our mechanistic studies suggested that RelB:p52 transcriptionally upregulated the expression of Axin1, which tethers β-catenin to its destruction complex. Inactivation of non-canonical NF-κB signaling reinforced the β-catenin-mediated expression of Raldh2, which promotes tolerogenic RA synthesis by DCs^22^. Ablating *Relb* or *Nfkb2* in DCs not only increased the frequency of intestinal Tregs but also improved the abundance of colonic IgA^+^ B cells, which fostered luminal IgA and the gut microbiome. Finally, haploinsufficiency of β-catenin in DCs lacking non-canonical NF-κB signaling reinstated colitogenic sensitivity in the composite knockout mice. Taken together, we chart a novel crosstalk between the non-canonical NF-κB pathway and the tolerogenic β-catenin-Raldh2 axis in DCs in tuning intestinal inflammation.

## Results

### Experimental colitis in mice is associated with and exacerbated by non-canonical *Relb*-*Nfkb2* signaling in DCs

To decipher if intestinal pathologies are linked to non-canonical NF-κB signaling in DCs, we interrogated single-cell RNA-seq data generated using colon tissues from mice subjected to dextran sodium sulfate (DSS)-induced acute colitis^23^ (Figure S1A). Based on the expression of prototypic macrophage- and DC-specific genes (Supplementary table S1), we categorized the intestinal mononuclear phagocyte (MNPs) population described in this dataset into macrophage and DC subsets (Figure 1A). Our subsequent analyses identified *Relb* or *Nfkb2* mRNA-expressing DCs in the gut of untreated mice (Figure 1B-1C). DSS treatment led to a rise in the abundance of mRNAs encoding these non-canonical NF-κB factors in intestinal DCs at day 6. We also observed an increase in the frequency of DCs expressing these mRNAs in the colitogenic gut (Figure 1C). Based on a previously described bulk transcriptomics study involving RelB-deficient mouse DCs^16^, we then deduced a panel of five top RelB-important genes - *Fgr*, *Top1*, *Crk*, *Gpd2* and *Cript* - that were also less reliant on other NF-κB factors for their expressions. We determined the abundance of corresponding mRNAs as a surrogate of non-canonical NF-κB activity. We found DCs from mouse colon possessed detectable levels of *Fgr*, *Top1*, and *Cript* transcripts even prior to DSS treatment (Figure 1D). Except for *Top1*, DSS treatment also led to elevated expressions of all other genes in intestinal DCs that were accompanied by an increased frequency of DCs expressing these mRNAs (Figure 1D). Our UMAP also captured the appearance of a distinct DC-population in the intestine of DSS-treated mice. Cataloguing DCs into classical DC1(cDC1) and cDC2 clarified that this population consisted of cDC2 (Figure S1B). However, we didn’t see discernable difference between these DC subsets with respect to the non-canonical NF-kB pathway engagement (Figure S1C). These studies implied DC-intrinsic non-canonical NF-κB signaling in experimental colitis. To demonstrate non-canonical NF-κB activation in intestinal DCs directly, we harvested DCs from gut-draining mesenteric lymph nodes (MLNs) as CD11c^+^CD64^-^MHCII^high^ cells and performed immunoblot analyses. Intestinal DCs from untreated mice displayed basal processing of p100 into p52, a hallmark of non-canonical NF-κB signaling (Figure 1E, S1D). DSS treatment further augmented p100 processing in these DCs in mice. We concluded that intestinal DCs possessed a basally active non-canonical *Relb*-*Nfkb2* pathway and that aberrant intestinal inflammation was associated with intensified non-canonical NF-κB signaling in these cells.

**Figure 1:**
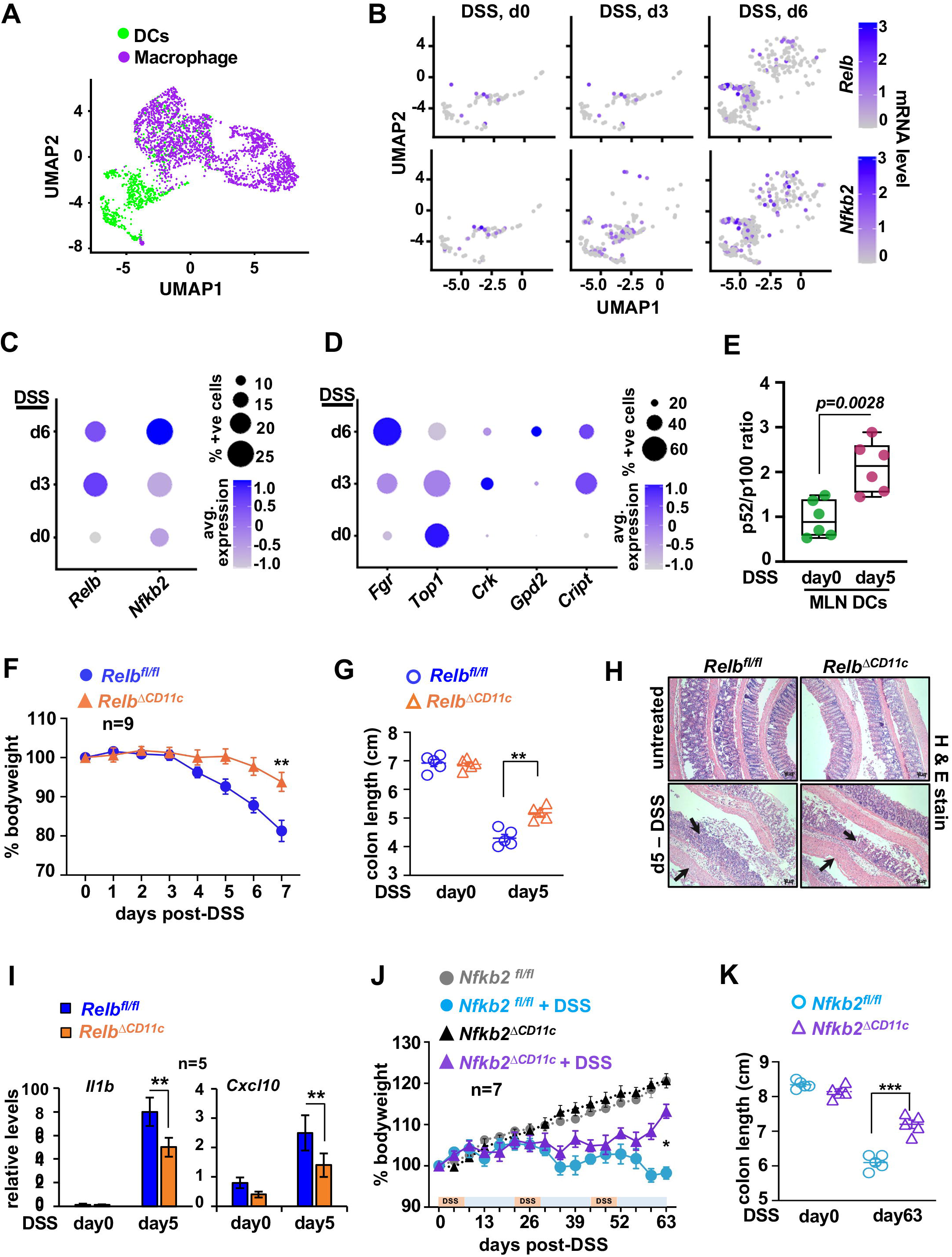
Charting the role of the non-canonical NF-κB pathway in DCs in modulating experimental colitis in mice. **A.** UMAP depicting the DC and macrophage cell population among mononuclear phagocytes in the mouse colon. Single-cell RNA-seq data available at the NCBI database was used for analyses (Ref: GSE148794). **B.** UMAP depicting the expression of *Relb* and *Nfkb2* mRNAs in a time course in intestinal DCs from WT mice subjected to DSS-induced acute colitis. **C.** and **D.** Dot plot revealing quantified levels of indicated mRNAs in intestinal DCs from the WT mice left untreated or subjected to DSS treatment. **E.** Boxplot (left) showing the abundance of p52 in relation to p100 in intestinal DCs, as determined by quantifying the respective band intensities in immunoblot analyses. DCs were collected from MLNs of untreated mice or those administered with 2% DSS in drinking water for 5 days. **F.** and **G.** *Relb^fl/fl^* and *Relb^ΔCD^*^11c^ mice were subjected to acute DSS treatment for 7 days and changes in the bodyweight **(F)** and colon length **(G)** were monitored. **G.** RT-qPCR demonstrating the colonic abundance of indicated mRNAs in untreated or DSS-treated *Relb^fl/fl^* and *Relb^ΔCD^*^11c^ mice. **H.** Representative images of H&E-stained colon sections from the indicated mice left untreated or subjected to acute DSS treatment. The data represent n = 5; 4 fields per section and a total of 5 sections from each set were examined. Black arrows indicate edema, leucocyte infiltration and epithelial lining damage. Scale bar, 50μm. **I.** Scatter plot revealing the serum concentration of FITC-dextran in *Relb^fl/fl^* and *Relb^ΔCD^*^11c^ mice left untreated or subjected to acute DSS treatment; FITC-dextran was gavaged orally 6h prior to serum collection. **J.** RT-qPCR demonstrating the colonic abundance of indicated mRNAs in untreated or DSS-treated *Relb^fl/fl^* and *Relb^ΔCD^*^11c^ mice. **K.** and **G.** In a chronic colitis regime, *Nfkb2^fl/fl^* and *Nfkb2^ΔCD^*^11c^ mice were subjected to 3 cycles of DSS treatment, where each cycle involved 7 days of 1% DSS treatment followed by 14 days of recovery. Subsequently, changes in the bodyweight **(F)** and colon length **(G)** were measured. Data represents mean ± SEM. For statistical analyses, two-tailed Student’s t-test was performed. *P < 0.05; **P < 0.01; ***P < 0.001; ns, not significant.

Next, we asked if the non-canonical NF-κB pathway in DCs caused colitogenic pathologies. Therefore, we generated *Relb^ΔCD^*^11c^ mice with DC-specific RelB deficiency by crossbreeding *Relb^fl/fl^* mice with CD11c-Cre^+^ mice (Figure S1E) and subjected these mice to experimental colitis. As such, acute DSS challenge for 7 days led to up to 20% bodyweight loss in littermate *Relb^fl/fl^*controls, accompanied by severe colon shortening, and pervasive epithelial disruption associated with edema and leukocyte infiltration (Figure 1F-1H, S1F). *Relb^ΔCD^*^11c^ mice presented significantly less reduction in bodyweight over the course of DSS treatment and a modest decrease in colon length with a relatively intact epithelium at day 5 (Figure 1F-1H). The intestinal epithelial architecture of untreated *Relb^fl/fl^* and *Relb^ΔCD^*^11c^ mice was histologically indistinguishable. RelB deficiency in DCs also restricted the DSS-induced increase in barrier permeability in knockout mice (Figure S1G). Similarly, our gene expression analyses revealed diminished DSS-induced expressions of inflammatory genes, including those encoding IL-1β and CXCL10, in the colon of *Relb^ΔCD^*^11c^ mice (Figure 1I).

Therefore, DC-intrinsic RelB deficiency moderated inflammation and acute colitis in mice. The coordinated functioning of proteins encoded by *Relb* and *Nfkb2* transduces non-canonical NF-κB signals. Compared to *Nfkb2^fl/fl^*controls, however, *Nfkb2^ΔCD^*^11c^ mice displayed only subtly improved bodyweight phenotype upon acute DSS treatment (Figure S1I-S1J). Interestingly, cyclical challenges with a low dose of DSS in the chronic chemical colitis regime resulted in a significantly less bodyweight reduction and colon shortening in *Nfkb2^ΔCD^*^11c^ mice at the end of the third cycle on day 63 (Figure 1J-1K). It has been reported that an alternate RelB:p50 heterodimer functionally compensates, albeit partially, for the absence of RelB:p52 activity in *Nfkb2*-deficient cells^24^. We reasoned that the molecular redundancies between RelB NF-κB factors obscured the phenotypic penetrance of *Nfkb2^ΔCD^*^11c^ mice in the acute colitis regime and that repeated DSS exposure unmasked colitis-resilient phenotypes in these mice because of exaggerated DC-mediated immune reactions in the chronic regime^25^. Overall, we conclude that experimental colitis in mice strengthens in DC the non-canonical NF-κB pathway, which exacerbates inflammatory intestinal pathologies in mice.

### Non-canonical NF-κB signaling restrains tolerogenic Raldh2 activity in DCs

To mechanistically understand the immunomodulatory functions of non-canonical NF-κB signaling, we set out to examine bone marrow-derived dendritic cells (BMDCs) from WT or knockout mice (Figure S2A). Our culture condition produced ∼ 85% CD11c^+^ BMDCs by day 9 that were associated with increased processing of p100 into p52 and elevated nuclear RelB and p52 levels (Figure 2A, Figure S2B). Our RNA-seq studies demonstrated that ablation of this constitutive non-canonical NF-κB signaling in *Nfkb2^-/-^* BMDC modified global gene expressions basally (Figure S2C). RA metabolism emerged as one of the top-ranking DC-relevant biological pathways enriched among genes differentially expressed between WT and *Nfkb2^-/-^* BMDCs (Figure 2B). Concordantly, our gene set enrichment analysis (GSEA) revealed a significant enrichment of RA targets among genes whose basal expressions were augmented in *Nfkb2^-/-^*BMDCs, indicative of strengthening autocrine RA actions in knockouts (Figure 2C). Sequential action of retinol dehydrogenases (Rdh) and retinal dehydrogenases (Raldh) convert retinol to RA, which is catabolized by the Cyp26 family of enzymes^26^ (Figure 2D). Further probing our RNA-seq data for the expression of genes encoding these enzymes revealed a consistently elevated mRNA level of Raldh2 in *Nfkb2^-/-^* BMDCs (Figure 2E). These studies indicated a possible role of non-canonical NF-κB signaling in transcriptionally modulating the RA pathway.

**Figure 2:**
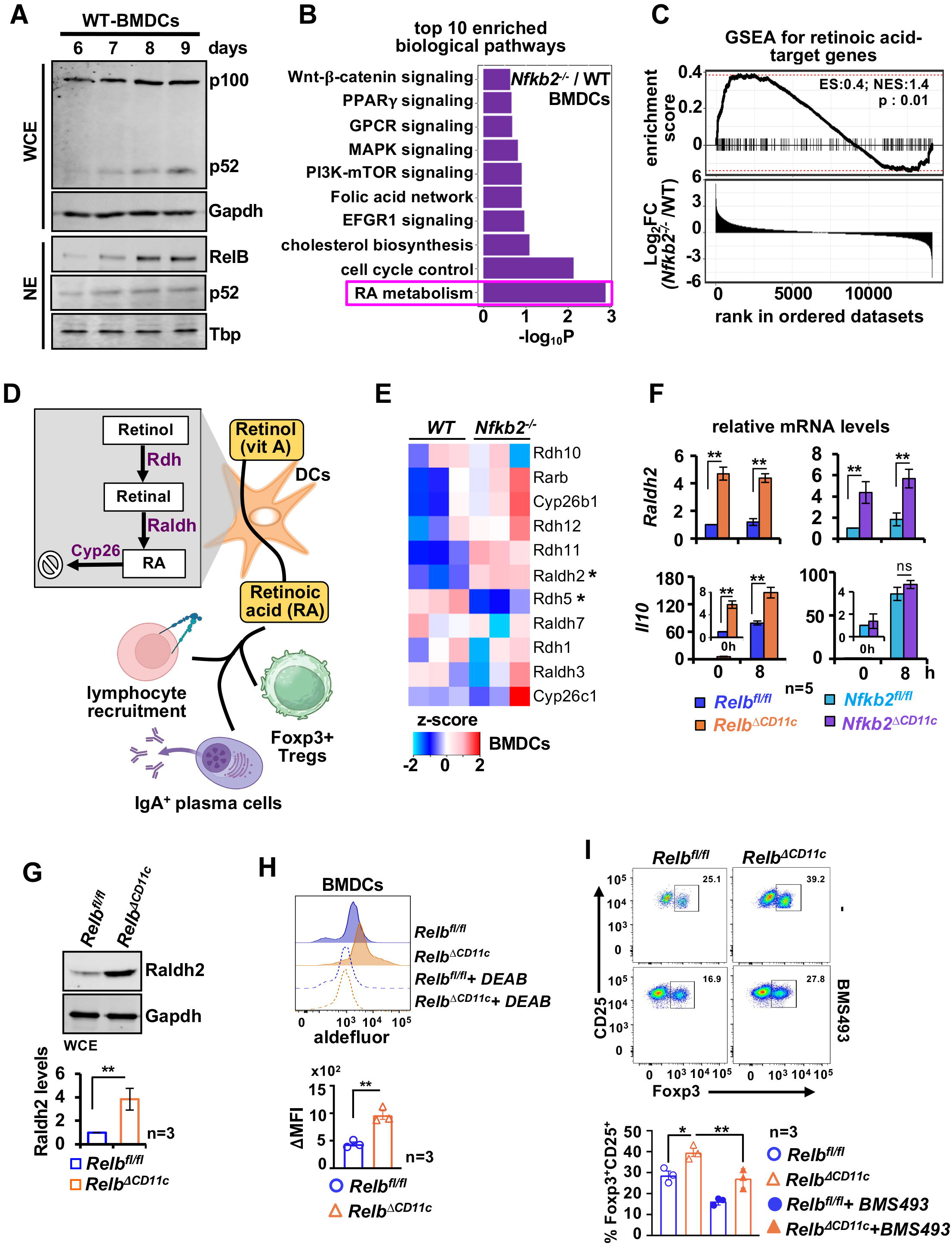
Defective *Relb* or *Nfkb2* functions enhance the expression and activity of Raldh2 in DCs. **A.** Immunoblot charting the abundance of p52 and p100 in whole cell extracts (top) or RelB and p52 in nuclear extracts (bottom) in cells harvested from BMDC-differentiating culture at the indicated days. **B.** Enrichment analyses revealing top 10 enriched DC-relevant biological pathways among genes differentially expressed between *Nfkb2^-/-^* and WT BMDCs. For each genotype, RNA-seq data from three independent biological replicates were used. **C.** GSEA comparing *Nfkb2^-/-^* and WT BMDCs for the enrichment of RA targets among differentially expressed genes (top). Fold change (FC) in the basal mRNA level between these genotypes was considered to create a ranked gene list, which was subjected to GSEA. Vertical lines represent individual RA targets. ES and NES denote enrichment and normalized enrichment scores, respectively. **D.** A cartoon describing the RA metabolism pathway and its functions in DCs. **E.** Heatmap showing the expression of selected RA metabolism-associated genes in WT and *Nfkb2^-/-^* BMDCs, as determined in RNA-seq. **F.** RT-qPCR comparing untreated or LPS-treated BMDCs derived from *Relb^fl/fl^* and *Relb^ΔCD^*^11c^ mice (left) or *Nfkb2^fl/fl^* and *Nfkb2^ΔCD^*^11c^ mice (right) for gene expressions. **G.** Representative immunoblot revealing the abundance of Raldh2 in *Relb^fl/fl^*and *Relb^ΔCD^*^11c^ BMDCs. Corresponding band intensities were quantified and presented as a barplot below. **H.** Histograms showing the Raldh enzymatic activity measured by Aldefluor assay in BMDCs. The Raldh inhibitor DEAB was used as a control. For a given genotype, ΔMFI was calculated as = MFIno DEAB - MFIDEAB, and presented in a barplot. **I.** Flow cytometry analyses demonstrating the generation of CD25^+^Foxp3^+^ CD4 Treg in DC-T cell co-culture *ex vivo*. Briefly, BMDCs were cultured with naïve T cells, isolated from the spleen of WT mice, in the presence of CD3 and TGF-β for 3.5 days. In specified sets, the RAR inhibitor BMS493 was added. The frequency of CD25^+^Foxp3^+^ Treg as a percentage of CD4 cells was determined and presented in the barplot (bottom). Data represent mean ± SEM. Two-tailed Student’s t-test was performed. *P < 0.05; **P < 0.01; ns, not significant.

To substantiate non-canonical NF-κB-mediated control of the RA pathway, we derived BMDCs from *Relb^ΔCD^*^11c^ or *Nfkb2^ΔCD^*^11c^ mice. As expected, *ex vivo* differentiated BMDCs from *Relb^ΔCD^*^11c^ or *Nfkb2^ΔCD^*^11c^ mice produced only a minor amount of RelB or p100, respectively (Figure S2D-S2E). In comparison to corresponding gene-sufficient floxed controls, *Relb^ΔCD^*^11c^ and *Nkfb2^ΔCD^*^11c^ BMDCs both displayed a ∼ 4.5-fold increase in the basal expression of Raldh2 mRNA in our RT-qPCR analyses (Figure 2F). LPS treatment did not discernably alter Raldh2 mRNA levels in WT or knockouts. In contrast, basal and LPS-induced IL-10 mRNA expressions were heightened in *Relb^ΔCD^*^11c,^ but not *Nfkb2^ΔCD^*^11c^ BMDCs. Because of the shared colitis resilience phenotype of *Relb^ΔCD^*^11c^ and *Nkfb2^ΔCD^*^11c^ mice, we focused on Raldh2, whose expression was similarly affected upon ablation of either *Relb* or *Nfkb2* in BMDCs. We detected a 3.8-fold excess accumulation of Raldh2 protein in immunoblot analyses and a 2.2-fold increase in the basal Raldh enzymatic activity in aldefluor assay in *Relb^ΔCD^*^11c^ BMDCs (Figure 2G-2H). *Nfkb2^ΔCD^*^11c^ BMDCs revealed similar increases in the Raldh2 protein level and enzymatic activity (Figure S2F-2G). Previous studies involving DC-T cell co-culture established RA as an important determinant of LP DC-mediated conversion of naïve T cells into Tregs *ex vivo* (Sun et al., 2007). Our own analyses revealed that *Relb^ΔCD^*^11c^ BMDCs were more proficient in converting naïve splenic T cells into CD25^+^Foxp3^+^ CD4 Tregs *ex vivo*, and BMS493, a retinoic acid receptor (RAR) antagonist, erased this RelB-effect in DC-T cell co-culture (Figure 2I). Therefore, DC differentiation *ex vivo* appears to trigger constitutive activation of non-canonical NF-κB signaling, which suppresses the tolerogenic Raldh2-RA axis.

### DC functions of the non-canonical NF-κB signal transducers limit the frequency of FoxP3^+^ Treg in the mouse colon

Next, we examined immunoregulatory attributes of non-canonical NF-κB-deficient DCs *in vivo*. Deletion of *Relb* or *Nfkb2* in CD11c^+^ cells did not significantly alter the frequency of DCs in MLNs or lamina propria (LP) in the colon (Figure 3A, Figure S3A). Also, when examined for the surface expression of CD80 and CD86 in flow cytometry analyses, intestinal DCs from *Relb^fl/fl^* and *Relb^ΔCD^*^11c^ mice were indistinguishable (Figure 3B). Similarly, *Nfkb2^fl/fl^* and *Nfkb2^ΔCD^*^11c^ mice displayed equivalent levels of these co-stimulatory molecules on intestinal DCs (Figure S3B). Corroborating BMDCs studies, intestinal DCs from *Relb^ΔCD^*^11c^ mice, but not from *Nfkb2^ΔCD^*^11c^ mice, expressed significantly more IL-10 (Figure 3B-3C). However, RelB-dependent IL-10 regulation was more apparent in MLN DCs. Importantly, we could confirm an increased level of Raldh2 mRNA in intestinal DCs from both *Relb^ΔCD^*^11c^ and *Nfkb2^ΔCD^*^11c^ mice (Figure 3D). Our enzymatic assay substantiated a more than 2.1-fold rise in the Raldh activity in intestinal DCs from untreated *Relb^ΔCD^*^11c^ mice (Figure 3E). Thus, non-canonical NF-κB signaling regulated Raldh2 activity in DCs *in vivo*.

**Figure 3:**
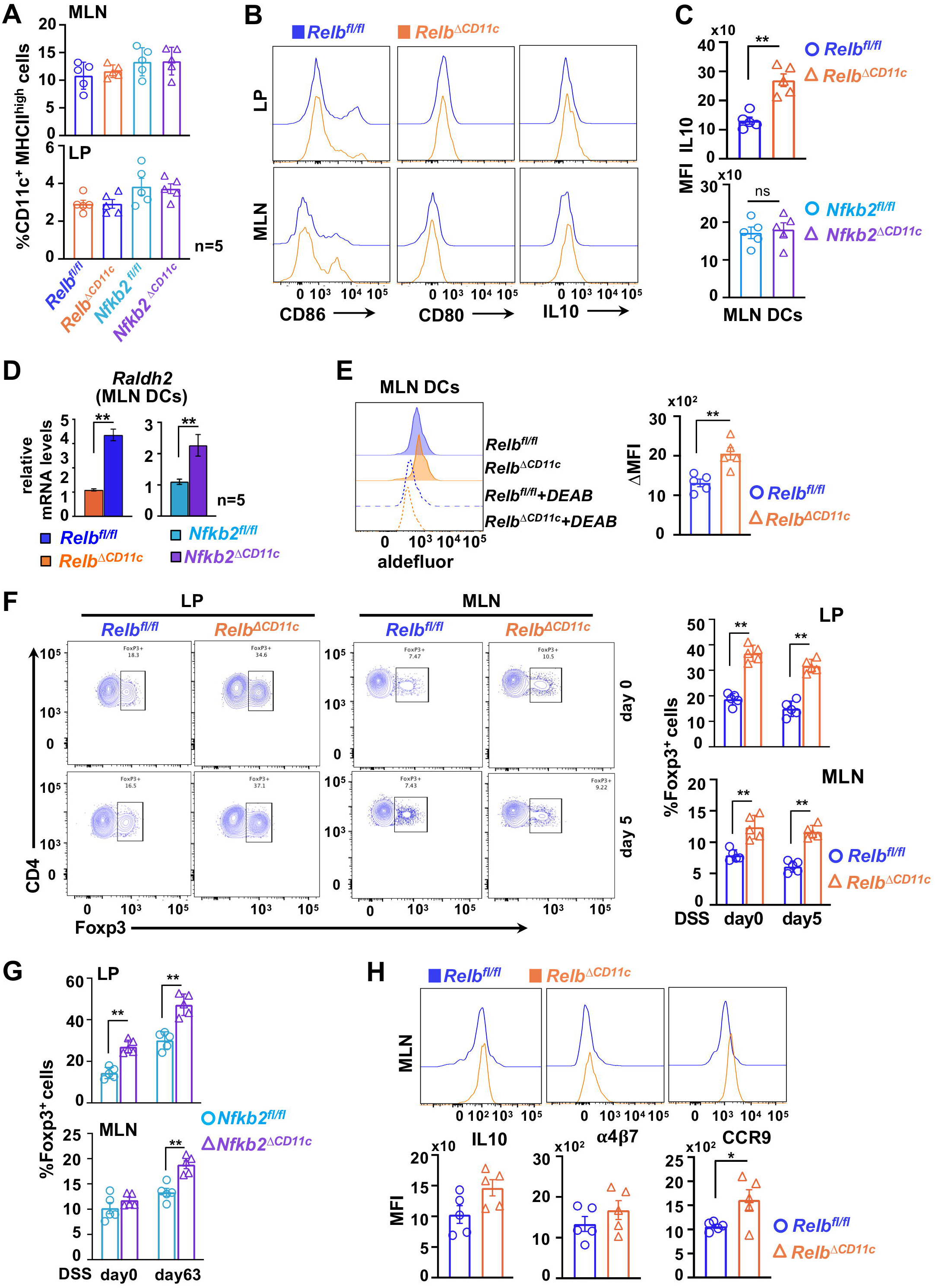
Disruption of non-canonical NF-κB signaling in CD11c^+^ cells accumulate FoxP3^+^ Treg in the mouse colon. **A.** Flow cytometry studies indicating the abundance of DCs, as a percentage of CD45+ cells, in MLN or LP. **B.** Representative histograms showing the expression of CD80, CD86 and IL-10 in DCs from LP or MLN from *Relb^fl/fl^* and *Relb^ΔCD^*^11c^ mice. **C.** Barplot revealing MFI for IL-10 expression in MLN DCs from the indicated mice. **D.** RT-qPCR depicting the abundance of Raldh2 mRNA in MLN DCs from *Relb^fl/fl^* and *Relb^ΔCD^*^11c^ or *Nfkb2^fl/fl^* and *Nfkb2^ΔCD^*^11c^ mice. **E.** Histograms capturing Raldh enzymatic activity in MLN DCs from *Relb^fl/fl^* and *Relb^ΔCD^*^11c^ mice. Quantified average MFI has been presented in a barplot. **F.** Representative contours from flow cytometry analyses showing the frequency of Foxp3^+^ Tregs as a percentage of CD4^+^ T cells in LP or MLNs of untreated or DSS-treated *Relb^fl/fl^* and *Relb^ΔCD^*^11c^ mice. Corresponding quantified data has been presented as a barplot in the right. **G.** Barplot revealing the mean frequency of Foxp3^+^ Treg in LP and MLN of *Nfkb2^fl/fl^* and *Nfkb2^ΔCD^*^11c^ mice. **H.** Overlapping histograms depicting the expression of IL-10, α4β7 and CCR9 in Foxp3^+^ Tregs from MLNs of *Relb^fl/fl^* and *Relb^ΔCD^*^11c^ mice. Quantified average MFIs are indicated in the individual barplot below. Each datapoint signifies an individual mouse; data represent mean ± SEM. Two-tailed Student’s t-test was performed. *P < 0.05; **P < 0.01; ***P < 0.001; ns, not significant.

Raldh2 in intestinal DCs drives the synthesis of RA, which supports FoxP3 expressions and upregulates gut-homing receptors α4β7 and CCR9 in Tregs^27,28^. Indeed, the FoxP3^+^ Treg frequency among CD4 T cells in LP and MLNs improved from 18.7% and 7.9% in *Relb^fl/fl^* mice to 36.6% and 12.4% in *Relb^ΔCD^*^11c^ mice, respectively (Figure 3F). While acute DSS treatment subtly skewed this Treg compartment at day 5, *Relb^ΔCD^*^11c^ mice maintained an overall increased level of these cells even in the colitogenic gut. Except for MLNs from untreated sets, the frequency of FoxP3^+^ Tregs was similarly augmented in the intestine of *Nfkb2^ΔCD^*^11c^ mice (Figure 3G, Figure S3C). Notably, FoxP3^+^ Tregs from MLNs of *Relb^ΔCD^*^11c^ mice were, if anything, subtly more efficient than those from *Relb^fl/fl^* counterparts in expressing IL-10 and α4β7, and displayed significantly elevated levels of CCR9 on their surface (Figure 3H). Inactivation of *Relb* or *Nfkb2* in CD11c^+^ cells did not discernably alter the frequency of total CD4 T lymphocytes in the colon of untreated or DSS-treated mice (Figure S3D). Together, non-canonical NF-κB signaling impedes the Raldh2-mediated RA synthesis pathway in mucosal DCs, limiting the abundance of Tregs, and their gut-homing receptor expressions, in the intestine.

### DC-intrinsic RelB function controls the abundance of IgA-secreting cells in the colon and gut microbiome

Previous studies suggested that RA promotes IgA-secreting cells at the gut mucosal interface^29^. We further explored whether reinforcing the RA pathway in non-canonical NF-κB-deficient DCs impacted intestinal sIgA levels. Our flow cytometry analyses disclosed a ∼ 2.5-fold increase in the frequency of IgA^+^B220^+^ cells in LP of *Relb^ΔCD^*^11c^ mice (Figure 4A-4B), and our ELISA ascertained a significantly elevated fecal sIgA levels in this knockout (Figure 4C). Because sIgA coating is known to shape the gut microbiome^5,30^, we performed 16s rRNA gene sequencing of fecal DNA to estimate the prevalence of different microbial taxa. We found that RelB deficiency was associated with changes specifically in two phyla; it led to an increase in the abundance of Firmicutes and a reduction in the level of Actinobacteriota (Figure 4D). Another prominent phylum Bacteroidota showed a comparable fecal abundance among *Relb^fl/fl^* and *Relb^ΔCD^*^11c^ mice. Genus-level analyses of sequencing data similarly revealed gut microbiome alterations in these mice, including an expansion of *Sangeribacter* and *Lactobacillus* but also a reduction of *Akkermansia* and *Faecalibacterium* abundance (Figure 4E). Corroborating DNA sequencing studies, our qPCR analyses of fecal DNA substantiated changes in Firmicutes and Actinobacteriota in *Relb^ΔCD^*^11c^ mice (Figure 4F). Moreover, MACS-based fractionation of fecal pellet identified enrichment of *Enterobacter*, *Bifidobacterium*, Segmented filamentous bacteria (SFB), *Sutterella* and *Prevotella* among IgA-bound gut microbes in *Relb^ΔCD^*^11c^ mice (Figure 4G). We also observed a decrease in the abundance of *β-proteobacteria* in the fecal IgA^+^ fraction in these knockouts.

**Figure 4:**
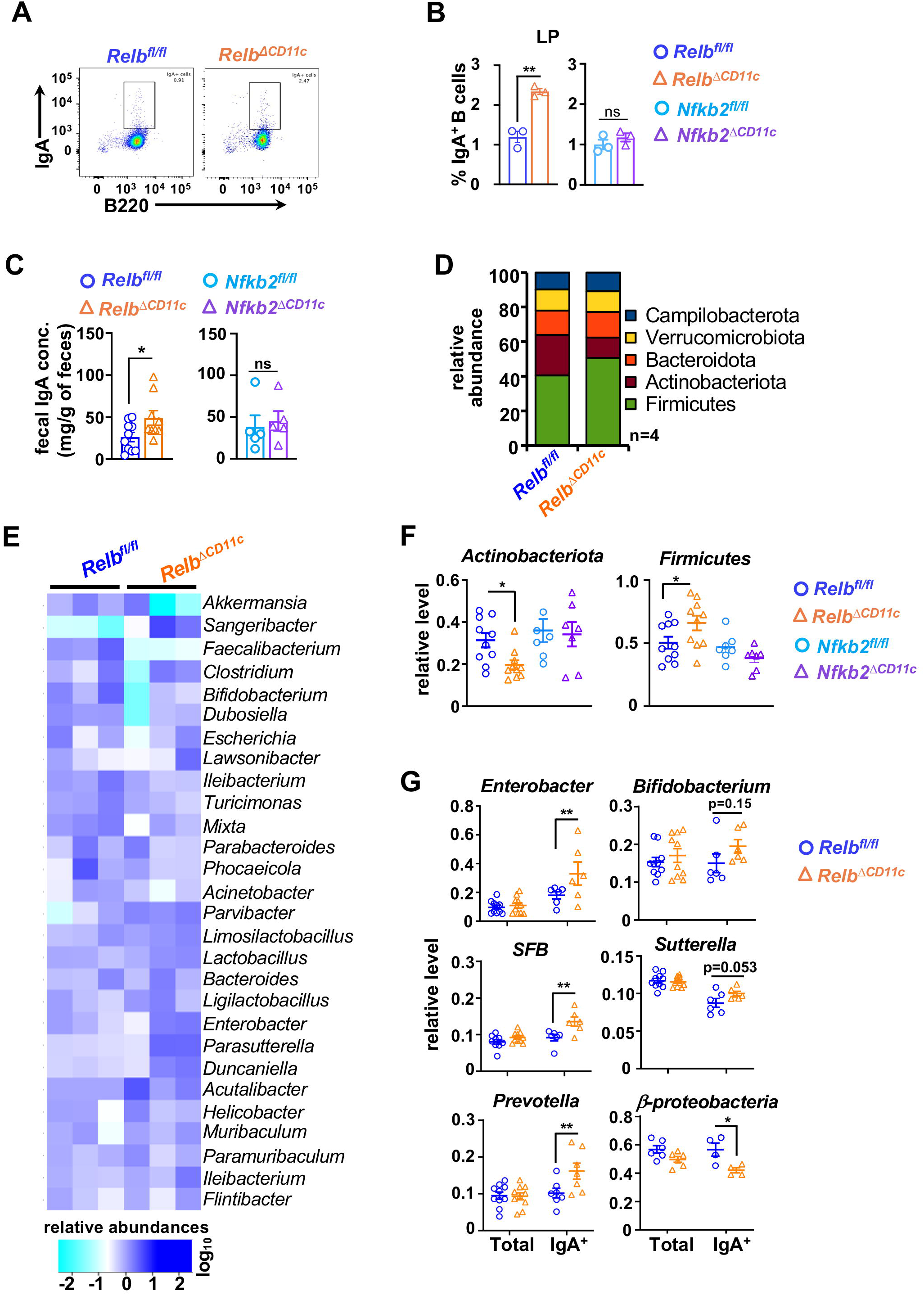
RelB-NF-κB deficiency in CD11c^+^ cells as a determinant of the intestinal abundance of IgA-secreting cells and gut microbiome. **A.** Representative FACS plots revealing the frequency of IgA^+^B220^+^ cells as a percentage of CD19^+^ cells in LP of *Relb^fl/fl^* and *Relb^ΔCD^*^11c^ mice. **B.** Barplot demonstrating quantified frequency of IgA^+^ B220^+^ cells in LP of mice of the indicated genotypes. **C.** ELISA comparing *Relb^fl/fl^* and *Relb^ΔCD^*^11c^ mice for sIgA concentrations in fecal pellets. **D.** and **E.** The relative abundance of various microbial phyla **(D)** and genera **(E)** charted in barplot **(D)** and heatmap **(E)**, respectively. Briefly, 16s rDNA sequencing was performed using fecal DNA from *Relb^fl/fl^* and *Relb^ΔCD^*^11c^ mice and prevalent phyla and genera were plotted. **F.** 16s rDNA-based qPCR analyses of fecal DNA showing the level of indicated microbial taxa in *Relb^fl/fl^* and *Relb^ΔCD^*^11c^ or *Nfkb2^fl/fl^* and *Nfkb2^ΔCD^*^11c^ mice. **G.** qPCR analyses MACS-sorted IgA^+^ fractions of fecal pellets comparing *Relb^fl/fl^* and *Relb^ΔCD^*^11c^ mice for the prevalence of the indicated bacteria. Each datapoint represents an individual mouse; data represent mean ± SEM. Two-tailed Student’s t-test was performed for Figure 5A-5C and one-way ANOVA was carried out for Figure 5F-5G. *P < 0.05; **P < 0.01; ns, not significant.

Notably, Firmicutes, and more so the increased ratio of Firmicutes to Bacteroidota, have been linked to improved intestinal barrier integrity^31,32^. Likewise, *Sangeribacter*, *Lactobacillus*, and *Faecalibacterium* are thought to largely promote intestinal health^33^. Therefore, barring a reduction in *Faecalibacterium*, *Relb^ΔCD^*^11c^ mice had an overall preponderance of gut-beneficial taxa. More so, these changes were associated with differential sIgA coating of microbial entities. Importantly, differential sIgA-coating not only quantitatively impacts the microbiome composition but also qualitatively affects gene expressions and metabolic functions of microbial entities in the intestinal niche^5^ . Curiously, *Nfkb2^fl/fl^*and *Nfkb2^ΔCD^*^11c^ mice were not discernibly different with respect to the frequency of intestinal IgA^+^B220^+^ cells, luminal sIgA levels and the abundance of Firmicutes and Actinobacteriota (Figure 4B-4C, Figure 4F). Taken together, we deduce that a DC-intrinsic RelB function, where *Nfkb2* is dispensable, limits the accumulation of intestinal IgA^+^ B cells and luminal IgA, which likely foster a gut microbiome protective against experimental colitis in mice.

### Non-canonical NF-κB signaling limits β-catenin-driven Raldh2 synthesis in DCs by directing Axin1 transcription

It was earlier reported that Notch2, Irf4, and β-catenin as key factors directing the transcription of *Raldh2* in DCs^22,34,35^. We found that RelB depletion increased the abundance of specifically β-catenin, and not Notch2 or Irf4, in BMDCs (Figure 5A). Compared to respective controls, total β-catenin levels were elevated 4.2-fold in *Relb^ΔCD^*^11c^ and 3.6-fold in *Nfkb2^ΔCD^*^11c^ BMDCs (Figure 5B). This was accompanied by a proportionate increase in the level of active β-catenin and nuclear β-catenin in these knockouts. Importantly, pharmacological inhibition of β-catenin-mediated transcription in non-canonical NF-κB-deficient BMDCs using iCRT3 restored the Raldh2 protein abundance to that observed in control cells (Figure 5C). Therefore, our data supported a role of β-catenin in boosting Raldh2 production in the absence of non-canonical NF-κB signaling.

**Figure 5:**
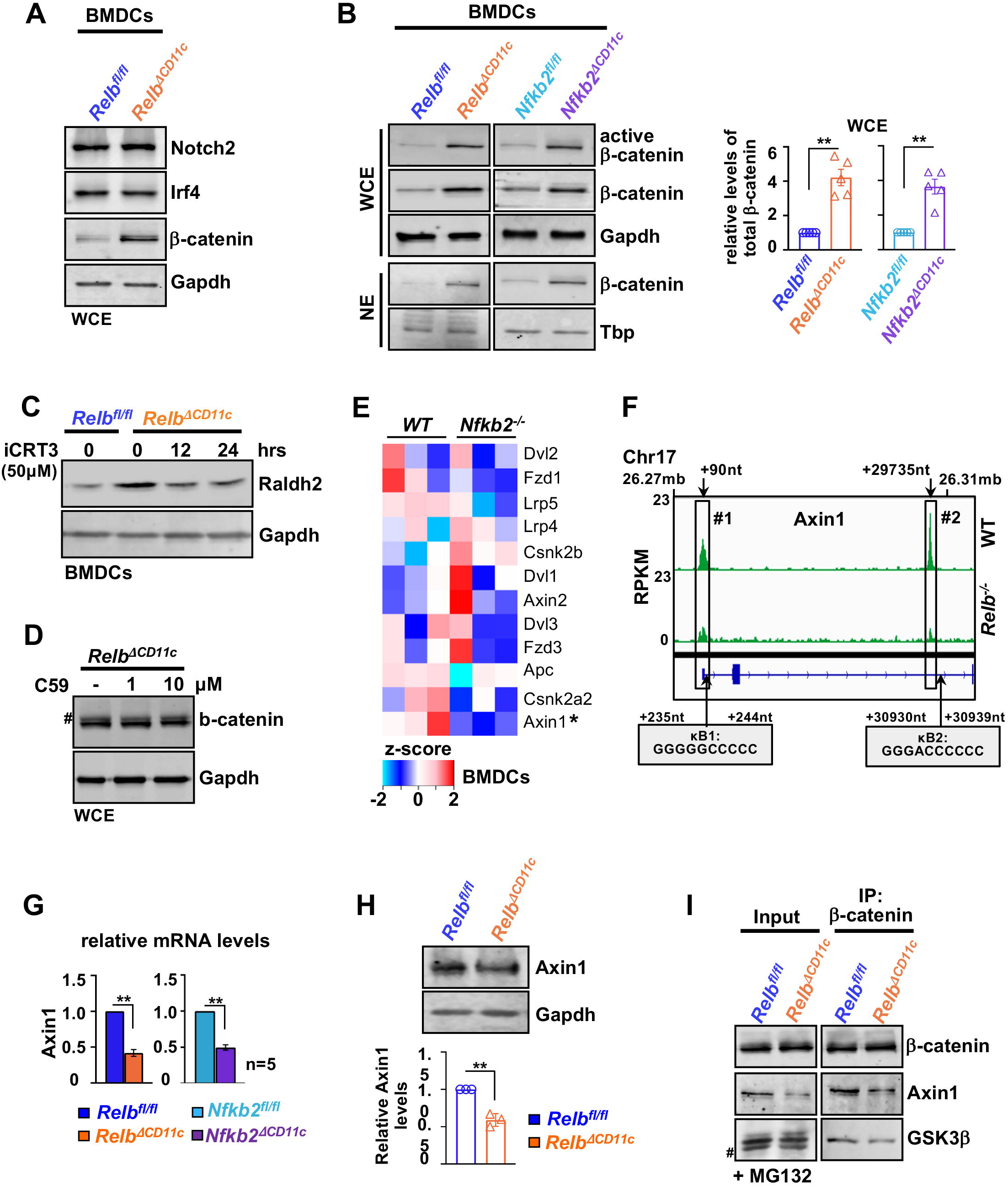
Reduced transcription of Axin1 promotes β-catenin-driven Raldh2 synthesis in noncanonical NF-κB-deficient DCs. **A.** Immunoblot analyses comparing *Relb^fl/fl^* and *Relb^ΔCD^*^11c^ BMDCs for the abundance of the indicated transcription factors. **B.** Representative immunoblot revealing active and total β-catenin levels in whole-cell or nuclear extracts derived from BMDCs of the indicated genotypes. Total β-catenin band intensities were also quantified and presented as barplot (right). Gapdh and Tbp served as loading controls for whole-cell and nuclear extracts, respectively. **C.** Immunoblot charting Raldh2 mRNA level in *Relb^fl/fl^* and *Relb^ΔCD^*^11c^ BMDCs left untreated or treated with 50μM of iCRT3, which inhibits β-catenin-mediated transcription, for 12h. **D.** Immunoblot analyses capturing the abundance of β-catenin in *Relb^ΔCD^*^11c^ BMDCs treated for 24h with 10μM of C59, which prevents secretion of Wnt ligands by inhibiting Porcupine. **E.** Heatmap comparing the abundance of mRNAs encoding the indicated β-catenin regulators in WT and *Nfkb2^-/-^* BMDCs, as determined by RNA-seq. **F.** The genome browser track of RelB ChIP-seq peaks in WT BMDCs (GSE236531) for *Axin1*. Cognate κB motifs associated with these peaks have also been indicated. **G.** and **H.** RT-qPCR **(G)** and immunoblot **(H)** analyses scoring the level of Axin1 mRNA **(G)** and protein **(H)**, respectively, in BMDCs. For immunoblot analyses, corresponding band intensities were quantified and presented (bottom). **I.** Immunoblot analyses of β-catenin co-immunoprecipitants obtained using *Relb^fl/fl^* and *Relb^ΔCD^*^11c^ BMDCs subjected to MG-132 treatment for 4h. Left, MG-132-treated cell extracts used as inputs for immunoprecipitation. MG-132 treatment was carried out to achieve comparable β-catenin levels between WT and knockout BMDCs. # represents non-specific bands. Quantified data represent the mean of at least three experimental replicates ± SEM. Two-tailed Student’s t-test was performed. *P < 0.05; **P < 0.01.

As such, Axin proteins tether β-catenin to a multiprotein destruction complex consisting of adenomatous polyposis coli (APC) and glycogen synthase kinase-3 (GSK-3) that promotes phosphorylation-dependent proteasomal degradation of β-catenin in resting cells^36^ (Figure S4A). Wnt ligands signal through Frizzled receptors to rescue -catenin from this constitutive degradation. We argued that strengthening autocrine signaling owing to increased Wnt-secretion augmented β-catenin levels in knockouts. However, blocking the autocrine pathway with C59, a small molecule inhibitor of Wnt secretion from cells, did not discernibly change the abundance of β-catenin in *Relb^ΔCD^*^11c^ BMDCs (Figure 5D). We then investigated if non-canonical NF-κB deficiency instead impacted other β-catenin regulators. Examining our RNA-seq data indeed identified a significant decline in the mRNA level of the Axin1 isoform in *Nfkb2^-/-^* BMDCs (Figure 5E). When we analyzed the published RelB ChIP-seq dataset generated using WT BMDCs, the genome browser track of *Axin1* revealed two distinct RelB peaks associated with a promoter and an intronic enhancer-κB element (Figure 5F). Our RT-qPCR studies substantiated that a dysfunctional non-canonical NF-κB pathway caused elevated expressions of Axin1 mRNA and protein in *Relb^ΔCD^*^11c^ and *Nfkb2^ΔCD^*^11c^ BMDCs (Figure 5G-5H). Notably, our immunoprecipitation experiments demonstrated that depleting Axin1 levels in *Relb^ΔCD^*^11c^ BMDCs impeded the association of β-catenin with GSK3β, a critical component of the destruction complex (Figure 5I). We conclude that non-canonical NF-κB-mediated transcriptional control of Axin1 suppresses β-catenin-driven Raldh2 expressions in DCs.

### β-catenin determines immunoregulatory DC functions of the non-canonical NF-κB pathway in the mouse gut

Because DC-intrinsic β-catenin functions were shown to limit mucosal inflammation in the intestine^22,37^, we asked if β-catenin accumulated in non-canonical NF-κB-deficient DCs was responsible for the resilience of these knockout mice to colitogenic insults. Corroborating our *ex vivo* analyses, we observed a ∼ 3-fold increase in the abundance of β-catenin in intestinal DCs from *Relb^ΔCD^*^11c^ or *Nfkb2^ΔCD^*^11c^ mice compared to those from respective control floxed mice (Figure 6A, Figure S4B). To causally establish the role of β-catenin in non-canonical NF-κB-deficient DCs, we then introduced a DC-specific β-catenin haploinsufficiency in mice devoid of RelB expression in DCs. Convincingly, MLN DCs from *Ctnnb1^fl/+^Relb^ΔCD^*^11c^ mice displayed a significant drop in the Raldh2 activity from those observed in *Relb^ΔCD^*^11c^ mice (Figure 6B). Consistent with the tolerogenic role of DC-expressed Raldh2, the frequency of colonic FoxP3^+^ Tregs and fecal sIgA in *Ctnnb1^fl/+^Relb^ΔCD^*^11c^ mice were restored to those found in *Relb^fl/fl^* mice (Figure 6C-6D). When challenged in the acute regime, *Ctnnb1^fl/+^Relb^ΔCD^*^11c^ mice exhibited bodyweight loss and colon shortening at day 6, which were rather analogous to *Relb^fl/fl^* mice (Figure 6E-6F). These studies illustrate a non-canonical NF-κB-driven DC network that aggravates intestinal inflammation by subverting the tolerogenic β-catenin-Raldh2-RA axis.

**Figure 6:**
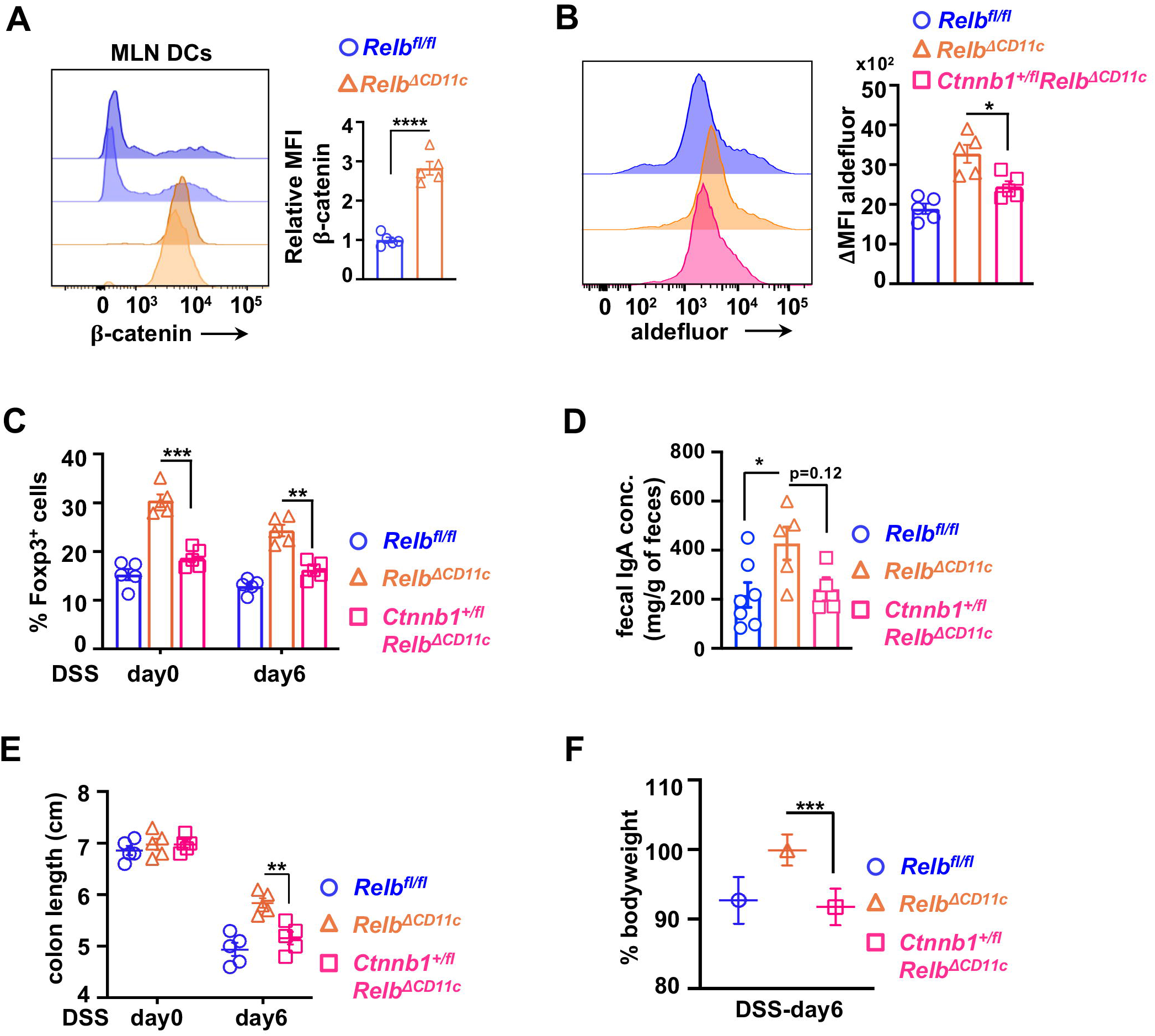
Haploinsufficiency of β-catenin in non-canonical NF-κB deficient DCs reinstates the colitogenic phenotype in mice. **A.** Histograms comparing *Relb^fl/fl^* and *Relb^ΔCD^*^11c^ mice for the expression of β-catenin in MLN DCs. Quantified MFI for -catenin has also been presented in a barplot. **B.** Histograms revealing the Raldh activity in MLN DCs from *Relb^fl/fl^*, *Relb^ΔCD^*^11c^ and *Ctnnb1^hetCD^*^11c^*Relb^ΔCD^*^11c^ mice. Corresponding ΔMFI values calculated from corresponding DEAB controls have been presented in the barplot. **C.** Barplot revealing the mean frequency of Foxp3^+^ Tregs in LP of indicated mice. **D.** ELISA comparing fecal IgA levels of the indicated genotypes. **E.** and **F.** Scatter plots revealing colon length and bodyweight changes on the day 6 upon acute DSS treatment of indicated mice. Each datapoint represents an individual mouse; data represent mean ± SEM. For statistical tests unless mentioned otherwise, One-way ANOVA was performed. **P < 0.01; ***P < 0.001, ****P < 0.0001; ns, not significant.

### Human IBD is associated with strengthening non-canonical NF-κB signaling in intestinal DCs

Finally, we sought to examine the relevance of our newly identified DC network in human IBD. To this end, we analyzed single-cell RNA-seq data generated using colon biopsies from 18 IBD patients and 12 healthy individuals^38^. Compared to healthy controls, intestinal DCs in the inflamed tissue of IBD patients possessed a substantially elevated level of *RELB* and *NFKB2* mRNAs (Figure 7A-7C). Although less apparent, DCs from the non-inflamed tissue of IBD patients also revealed an increase in the abundance of these mRNAs. We then determined the expression of RelB-important genes, viz *Fgr*, *Top1*, and *Crk*, in intestinal DCs from human subjects. Examining single-cell data revealed augmented DC levels of *FGR*, *TOP1*, *CRK* and *CRIPT* mRNAs in the inflamed as well as non-inflamed tissues from IBD patients that were associated with an increased frequency of DCs expressing these mRNAs in colonic tissues (Figure 7D). Therefore, our analyses suggested that human IBD was associated with heightened non-canonical NF-κB signaling in intestinal DCs.

**Figure 7:**
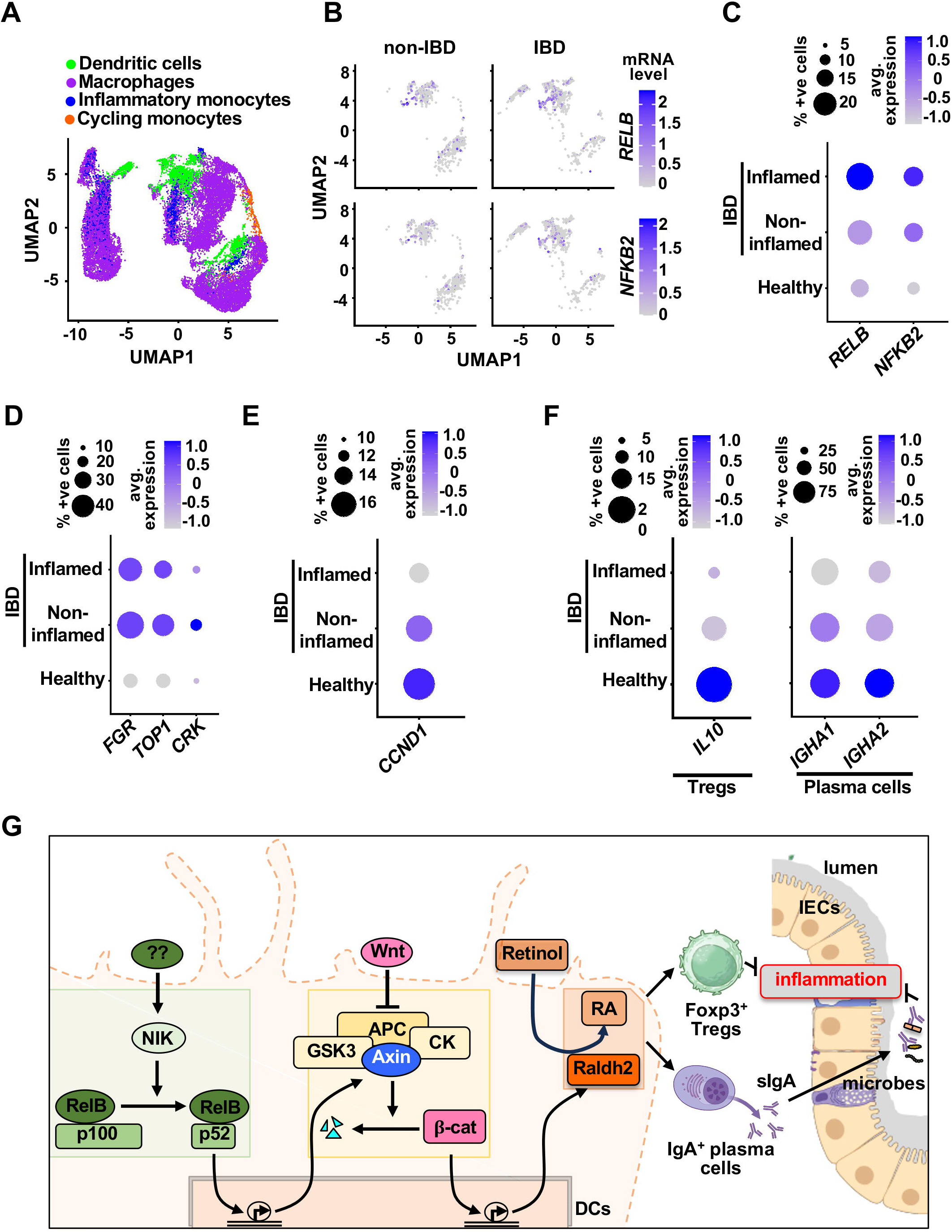
Engagement of non-canonical NF-κB-driven DC circuitry in human IBD. **A.** Feature plot depicting DCs among indicated myeloid cells in colonic tissues from IBD patients and healthy individuals. Briefly, single-cell RNA-seq data available at Single Cell Portal was used for analyses (Ref: SCP 259). **B.** UMAP comparing IBD patients and non-IBD subjects for the level of *RELB* and *NFKB2* mRNAs in intestinal DCs; the intensity of dot color denotes the abundance of corresponding mRNAs in each cell. **C.** Dot plot showing the corresponding expression levels of the indicated genes in DCs as depicted in panel B. The color gradient of dots represents the average expression level, while the dot size denotes the percentage of cells expressing a given gene. **D** - **E.** Dot plot illustrating the expression of indicated transcripts in intestinal DCs. **F.** Dot plot revealing the expression of *IL10* in Tregs (left), and *IGHA1* and *IGHA2* in plasma cells (right) from the colon of IBD patients and healthy subjects. **G.** A cartoon depicting a noncanonical NF-κB pathway-driven signaling circuitry in DCs that tunes intestinal inflammation.

Next, we probed if, corroborating our observation in knockout mice, strengthening non-canonical NF-κB signaling in DCs weakened the β-catenin-Raldh2-RA axis in IBD patients. While previous studies linked IBD to decreased Raldh activity in intestinal DCs^6^, we examined the available single-cell data^38^ for the expression of known β-catenin-regulated genes as a proxy for β-catenin’s transcriptional activity. Indeed, we could demonstrate that *CCND1*, a well-recognized β-catenin-activated gene, was expressed by approximately 20% of intestinal DCs and substantially downmodulated in IBD patients (Figure 7E). IBD was associated with a substantial decrease in the frequency of IL-10-expressing Tregs (Figure 7F, left) as also observed earlier^39^. We further observed a reduction in the average expression of *IGHA1* and *IGHA2,* genes encoding IgA heavy chain, in colonic plasma cells of the IBD patients compared to the non-IBD subjects (Figure 7F, right). Collectively, our investigation argues that the non-canonical NF-κB pathway targets β-catenin to limit the tolerogenic Raldh2 functions in intestinal DCs, orchestrating local inflammation in colitogenic mice as well as in IBD patients (Figure 7G).

## Discussion

Despite the known immunoregulatory role of the non-canonical NF-κB pathway in DCs^16–18,20^, whether RelB:p52 distorts DC functions in intestinal pathologies remains unclear. Our investigation identified intensified non-canonical NF-κB signaling in intestinal DCs from colitogenic mice and causally linked this pathway to exacerbated experimental colitis (Figure 1). DC-derived RA imparts tolerogenic attributes, and β-catenin induces the expression of the RA-synthesizing enzyme Raldh2 in intestinal DCs^22,27,40,41^. Our mechanistic studies suggested that RelB:p52 transcriptionally supported Axin1, which restrained Raldh2 synthesis in DCs by downmodulating β-catenin (Figure 2, Figure 5). Concordantly, non-canonical NF-κB impairment reinforced β-catenin-mediated Raldh2 expressions in DCs, improving the colonic frequency of protective Tregs and IgA^+^ B cells and also luminal sIgA levels, which were associated with a modified gut microbiome (Figure 3-4). Indeed, introducing β-catenin haploinsufficiency in RelB-deficient DCs reinstated the colitogenic sensitivity in mice (Figure 6). Corroborating our studies involving knockout mouse strains, the inflamed colon from IBD patients harbored DCs displaying heightened non-canonical NF-κB signaling and reduced Raldh activity, presumable owing to diminished β-catenin functions (Figure 7). Consistently, IBD patients also revealed a diminished frequency of FoxP3^+^ Tregs and IgA-secreting cells in the intestinal niche. IBD is multifactorial, with the involvement of several environmental, genetic, and immune components. Our analyses led us to propose DC functions of the non-canonical NF-κB pathway as a contributing factor to intestinal pathologies. In this mechanistic model, non-canonical NF-κB signaling subverts β-catenin-dependent, Raldh2-mediated synthesis of RA, which enforces a tolerogenic environment in the inflamed gut by promoting Tregs and luminal IgA (Figure 7H).

It was earlier reported that DC-specific ablation of *Relb*, but not *Nfkb2*, led to a systemic increase in the frequency of FoxP3^+^ Tregs^19^. We found a convergence of *Relb^ΔCD^*^11c^ and *Nfkb2^ΔCD^*^11c^ mice with respect to immune phenotypes – they both accumulated FoxP3^+^ Tregs in the intestine and were resilient to experimental colitis. Our data contended that systemic Treg effects were less important and that β-catenin-dependent modulation of specifically the intestinal Treg compartment was critical for the colitogenic role of RelB:p52 in DCs. As opposed to a substantial increase in the intestinal Treg frequency in *Relb^ΔCD^*^11c^ mice, RelB-depleted DCs *ex vivo* displayed only modest, albeit RA pathway-dependent, enhancement of Treg generation from naïve T cells. Based on the literature^28^ and our study, we reason that in addition to its role in priming Treg differentiation, DC-derived RA also promoted gut homing of Tregs and their stabilization in the intestinal niche in the knockout mice. Additionally, RelB in DCs was shown to promote microbiota-reactive RORγt+ Tregs in the small intestine^19^. Although our study did not distinguish between tissue- and microbiota-reactive Tregs in the colon, we speculate that the observed increase in the frequency of FoxP3^+^ Tregs in our knockouts could be driven by tissue Tregs. Importantly, other than microbiota-reactive Tregs, tissue Tregs were also found to be critical for tolerance to gut commensals^42^. Of note, it was suggested that NIK in DCs induced the expression of IL-23, which strengthened Th17 responses to *Citrobacter rodentium*^20^. A similar DC-intrinsic role of RelB in promoting IL-23-mediated Th17 response to gut commensals cannot be ruled out. However, our work argues that Th17 regulations by RelB in DC could be secondary non-canonical NF-κB-mediated Treg controls, at least in the intestinal niche.

We further noticed interesting differences between *Relb* and *Nfkb2* genotypes. In particular, *Nfkb2*-deficient DCs did not show elevated IL-10 expressions as observed in RelB-deficient DCs. *Relb^ΔCD^*^11c^ mice accumulated intestinal IgA^+^ B cells, whereas *Nfkb2^ΔCD^*^11c^ mice did not. Unlike *Relb^ΔCD^*^11c^ mice, *Nfkb2^ΔCD^*^11c^ mice did not readily produce a phenotype in the DSS-induced acute colitis regime. Previous studies suggested that IL-10, in addition to DC-derived RA, promotes IgA class switching in the gut^2^, and also documented an inhibitory role of RelB in DC-mediated IL-10 expressions^43^. Additionally, RelB:p50 heterodimer was shown to partially compensate for the lack of RelB:p52 in the *Nfkb2*-deficient cell system^24^. In light of these studies, we conjecture that RelB:p50 and RelB:p52 acted redundantly in DCs in downmodulating IL-10, while a non-redundant role of RelB:p52 in Axin1 transcription restrained β-catenin-dependent Raldh2 synthesis. Accordingly, ablating *Relb* in DCs supported the synergy between IL-10 and the RA pathway, culminating in the increased luminal IgA level. Conversely, RelB:p50 may have masked the phenotype of *Nfkb2^ΔCD^*^11c^ mice in the acute regime by preventing heightened IL-10 expressions in knockout DCs and, thereby, changes in the intestinal IgA^+^ cell compartment. *Nfkb2^ΔCD^*^11c^ mice also displayed a somewhat less marked increase in the frequency of intestinal Tregs compared to *Relb^ΔCD^*^11c^ mice, presumably owing to IL-10 expression differences between these genotypes. While the biochemical mechanism underlying RelB-mediated IL-10 control warrants further investigation, our comparison involving *Relb^ΔCD^*^11c^ and *Nfkb2^ΔCD^*^11c^ strains identified distinct as well as overlapping gene controls by the non-canonical NF-κB signal transducers *Relb* and *Nfkb2* in DCs with implications for intestinal health.

The widely known role of the non-canonical NF-κB pathway lies in lymphoid stromal cells in supporting the expression of homeostatic lymphokines, which promote secondary lymph node development during early embryogenesis and also lymphocyte ingress in lymph nodes in adults^15,44^. In addition, an *Nfkb2*-dependent cross regulatory mechanism was shown to aggravate colon inflammation by escalating RelA-driven inflammatory gene expressions in IECs^45^. Our work unequivocally established that non-canonical RelB:p52 NF-κB signaling in DCs inhibited the tolerogenic β-catenin-Raldh2 axis, fueling intestinal pathologies. Beyond tolerogenic DC functions, β-catenin and the Raldh2-RA pathway have other biological roles. For example, β-catenin deregulations in intestinal epithelial cells drive colon cancer^46^. In a recent study, Conlon et al (2020) suggested that LTβR inhibits WNT/β-catenin signaling in lung epithelial cells, exacerbating tissue damage in chronic obstructive pulmonary disease^47^. On the other hand, neuronal-derived RA promoted early lymph node biogenesis^48^, and intestinal DC-secreted RA was critical for the maintenance of secondary lymph nodes^49^. We argue that our work may provide for a significant revision of our understanding of non-canonical NF-κB functions in physiology and disease. To this end, future studies ought to examine the potential trigger of non-canonical NF-κB signaling in various anatomic niches and their plausible impact on β-catenin activity or the RA pathway in shaping immune responses or developmental processes.

Finally, micronutrients are emerging as important determinants of mucosal immunity and potential interventional tools for inflammatory ailments^50^. Previous studies indicated that vitamin D exerts a negative impact on RelB synthesis in DCs ^51,52^. As such, Raldh2 empowers intestinal DCs to secrete constitutively copious amounts of RA. Our current investigation posits that non-canonical NF-κB signaling curbs this immunomodulatory retinoic acid synthesis pathway in intestinal DCs, imparting vulnerability to colitogenic insults. Therefore, it appears that beyond its well-articulated role in immune gene expressions, DC-intrinsic non-canonical NF-κB signaling could be also critical for the micronutrient control of intestinal inflammation. While the significance of our study in the context of nutritional or DC-based therapy remains to be tested, we, in sum, establish a DC network integrating immune signaling and micronutrient metabolic pathways in the intestine.

## Acknowledgement

We thank Prof. Amitabha Bandyopadhyay, IIT Kanpur for help with *Ctnnb ^fl/fl^* mice. We thank V. Kumar for technical support and Dr. P. Nagarajan for the help with animal husbandry. Research in the PI’s laboratory was funded by DBT (BT/PR36631/BRB/10/ 1862/2020) and NII-Core. AD and NK thank DST-INSPIRE and DBT, respectively, for research fellowships.

## Author Contribution

A.D., N.K. and S.B. conceptualized the study and analyzed data. A.D. and N.K. performed the experiments with help from M.C., S.A.A., and B.C. Flow cytometry analyses were carried out with help from U.M. and guidance from A.A. 16s rRNA sequencing and data analyses were done with help from S.K. and supervision from B.D. Single-cell transcriptomics data was analyzed by N. K. and N.B. with guidance from D.S. ChIP-seq data analyses was performed by Sw.B. A.D., N.K. and S.B. wrote the manuscript.

## Material and methods

### Animal use

Sources of parental mouse strains used in this study have been indicated in the Key Resources Table. All strains were housed at the National Institute of Immunology and used in accordance with the guidelines of the institutional animal ethics committee (Approval no – IAEC – 487/18, 615/22, IBSC-512/22). CD11c-Cre mice were crossed with various floxed genotypes for generating CD11c cell-specific knockouts.

### Studying chemically-induced colitis in mice

Seven to eight weeks old littermate male mice of indicated genotypes were cohoused for at least one week prior to experiments. As described ^25^, acute colitis was induced by administering 1.5% DSS in drinking water for seven days. For chronic colitis induction, mice were fed with 1% DSS in drinking water for seven days followed by thirteen days of normal water; this cycle was repeated thrice. DSS-treated mice were examined for bodyweight changes in a time course. As a humane endpoint, mice with more than 20% bodyweight loss were immediately euthanized. For certain sets, mice were euthanized on indicated days post-DSS treatment and the length of the excised colon was measured. For histological studies, the colon was fixed in 4% formalin and embedded onto the paraffin block following the Swiss roll technique. Colon sections were examined following hematoxylin and eosin (H&E) staining or upon additional staining using Alcian Blue. Images were captured in an Olympus inverted microscope using Image-Pro6 software. To assess intestinal permeability, mice were gavaged with 440 mg/kg FITC-dextran 6h before fluorescence measurements in sera.

### Analyses of colonic immune cells

LP immune cells were isolated from dissected mouse colons, as described ^53^, except that an 80 and 40 percent Percoll gradient was performed and lymphocytes were collected from the interphase of Percoll layers. For MLNs and the spleen, single-cell suspensions were prepared by meshing these organs using a 70-micron cell strainer. Cells were then stained with fluorochrome-conjugated antibodies, and analyzed by flow cytometry. For surface staining, cells were labeled with antibodies in the staining buffer for 30 min on ice. Intracellular staining was performed using a commercially available kit as per manufacturer’s instruction (eBiosciences). For T cell analysis, cells were stimulated using leukocyte activation cocktail (BD Biosciences) for 4h before being subjected staining. Flow cytometry data were acquired on A5 symphony and analyzed using FlowJo v10 software. For sorting cells, BD FACSAria III cell sorter was used. Details of the gating strategy and flow cytometry antibodies have been provided in Supplementary Fig. 2 and the Supplementary Table S1, respectively.

### Studies involving BMDCs

BMDCs were generated from mouse bone marrow cells following a revised GM-CSF/IL-4 protocol that involves supplementing GM-CSF-containing DC-differentiation medium with rIL-4 but from day six of cell seeding^54^. Non-adherent cells were collected from the culture at day nine and seeded in fresh plates for experiments. BMDC differentiation was confirmed by examining the surface expression of CD11c and MHC-II by flow cytometry. Bone marrow cells from *Relb^ΔCD^*^11c^ and *Nfkb2^ΔCD^*^11c^ mice were used to produce BMDCs lacking the function of *Relb* and *Nfkb2*, respectively. Gene disruptions were confirmed in these day-nine BMDCs by immunoblot analyses.

For the Treg generation assay, naive CD4^+^ T cells (CD4^+^CD25^−^CD62L^+^) were sorted from the spleen of WT C57Bl/6 mice using a commercially available kit (Miltenyi Biotec). Next, 1×10^5^ naïve CD4^+^ T cells were co-cultured with WT or gene-deficient BMDCs at a ratio of 1:1 in the presence of a soluble anti-CD3 antibody (1μg/ml) and Treg polarizing cytokines - TGF-β (5ng/ml), rIL2 (20U/ml) and RAR inhibitor, BMS493 (100nM). After another three and a half days in the culture, cells were subjected to flow cytometry analyses for the expression of CD4, CD25, IFNγ, and FoxP3. RALDH activity was determined using the ALDEFLUOR assay kit, as per the manufacturer’s protocol (Stem Cell Technologies). Specifically, 0.5 million cells per ml were used for the analysis and incubated for 35 mins. The frequency of Aldefluor^+^ cells in the CD11c^+^MHCII^int/hi^ BMDC population was scored by flow cytometry.

### Biochemical and gene expression studies

Immunoblot analyses and RT-qPCR were performed essentially following our previously published methods ^45^ using WT or gene-deficient BMDCs. In certain experiments, BMDCs were also treated with 100ng/ml of LPS for 8h. For *in-vivo* analyses, cells were sorted from mice MLN using antibodies against required markers and similarly processed for immunoblotting. Primary antibodies and RT-qPCR primers used in this study have been described in the Supplementary Table S1 and Table S2. For Immunoblot assays, Cy5 conjugated secondary antibodies were used (Cytiva Lifesciences); gel images were acquired using Typhoon Variable Mode Imager, and band intensities were quantified in ImageQuant. For detecting active β-catenin, an antibody (CST, 8814S) that specifically recognizes unphosphorylated β-catenin simultaneously at Ser-33, Ser-37 and Thr-41 was used. As described earlier ^55^, whole-cell lysate prepared from ∼ 10^6^ cells was used for co-immunoprecipitation studies. Immunoprecipitation was performed using 2μg of anti-β-catenin antibody, and immunopellets were examined by immunoblot analyses using Trueblot secondary antibody. For colonic mRNA measurements, total RNA isolated from tissues derived from the distal colon was used.

For RNA-seq studies, BMDCs generated from WT and global *Nfkb2^-/-^*mice were compared in triplicates. Library preparation and sequencing were carried out at Nucleome Informatics, Hyderabad. When the cumulative read count of a transcript estimated from a total of six experimental sets was less than fifty, the corresponding gene was excluded from the analysis. Ensembl IDs lacking assigned gene names were also excluded. Accordingly, we arrived at a list of 19118 genes (available through GEO accession-GSE249101). Next, average read counts of these genes in experimental sets were calculated, and fold changes in the mRNA levels in *Nfkb2^-/-^* cells in relation to WT BMDCs were determined using the DEseq2 package from Bioconductor. Genes were then ranked in descending order of this fold change difference. Differentially expressed genes with fold change values above or below 1.2 were subjected to the pathway enrichment analysis using a list of Wikipathways available at MSigDB (www.gseamsigdb.org). The ranked gene list was also examined by GSEA involving the fgsea package using a previously described list of Retinoic acid targets ^56^. Integrative Genome viewer was used for generating genome browser tracks for RelB binding to the *Axin1* locus. For identifying NF-κB binding motifs around the ChIP-seq peaks, we used the Meme-Suite.

### Single cell RNA-Seq data analysis

We analyzed publicly-available single-cell RNA-Seq data generated using colonoscopic biopsies from human IBD patients (Single Cell Portal, SCP259)^57^, and those derived using colonic tissues from DSS-treated WT mice (GSE148794) ^58^. For human IBD, a total of 1819 colonic DCs from healthy or inflamed biopsies were examined. For DSS-induced colitis in mice, a total of 2236 intestinal mononuclear phagocytes (MNPs) from either untreated WT mice or those subjected to DSS treatment for 3 or 6 days were collectively considered. Leveraging a list of prototypic macrophage or DC genes, these MNPs were clustered into macrophages or DCs using the AUCell package in R. Then intestinal DCs were examined for the expression of indicated genes. Seurat package in R was used for gene expression analyses.

### Microbiome studies

Fresh feces were collected from eight weeks old littermate floxed and gene-deficient mice that were cohoused. Bacterial genomic DNA was extracted using QIAamp PowerFecal Pro DNA kit (Qiagen). The V3-V4 region of 16s rDNA was amplified by PCR and sequenced in the MiSeq platform at THSTI (Illumina) using the 2 × 250 bp paired-end protocol yielding pair-end reads that overlap almost completely. The read quality was examined by fastQC program (see Key Resource Table 3 for program source). Reads were processed based on a threshold phred score (Q phred) ≥20, and then merged to obtain high quality, long single reads using Paired-End reAd mergeR (PEAR) program (see Key Resource Table 3 for program source). The QIIME2 pipeline (qiime2-2022.2) was used for generating Operational Taxonomic Unit (OTU) table from the merged reads – reads were clustered based on 99% sequence identity. The naïve Bayes machine-learning classifier plugin of QIIME2’s q2-feature-classifier was used to find taxonomic classification up to the genus level based on SILVA SSU 138 (See Key Resource Table 3 for program source). The core OTU was subtracted from the main OTU table if an OTU was present in ≤80% of total sample of the group. The relative abundance of each OTU was calculated by number of the OTU in each sample divided by total number of OTUs in all the samples. The differential microbial abundance was converted into log 10 values to show negatively and positively abundant taxa in both the groups. The microbial alpha diversity analysis was performed using the ampvis2 R-package program. The statistical analysis was carried out using the tidyverse and ggplot2 R-packages (see Key Resource Table 3 for program source).

Further, the composition of bacterial taxa in flow cytometry-sorted IgA+ and IgA-fractions of the fecal microbiota was assessed using qPCR with the primers described in Key Resource Table 2. For measuring fecal IgA by ELISA, feces were homogenized in sterile PBS, and centrifuged to remove bacteria and insoluble debris. Fecal samples were then serially diluted and analyzed for IgA concentration by ELISA using a commercially available kit following manufacturer’s protocol (Invitrogen). Reading was acquired in a microplate reader (MultiSkan FC microplate photometer).

## Data and Software Availability

The MIAME version of the RNA seq dataset is available on NCBI Gene Expression Omnibus accession GSE249101, and the microbiome data is available at the accession PRJNA1046443.

## Supplemental Information

Supplemental Information consists of four supplementary figures, supplementary table S1, and Key resource tables1-3.

