## Supplementary material for "Non-canonical NF-κB signaling promotes intestinal inflammation by restraining the tolerogenic β-catenin-Raldh2 axis in dendritic cells": 07 Suppl Fig Deka and Kumar et al_Final.pdf

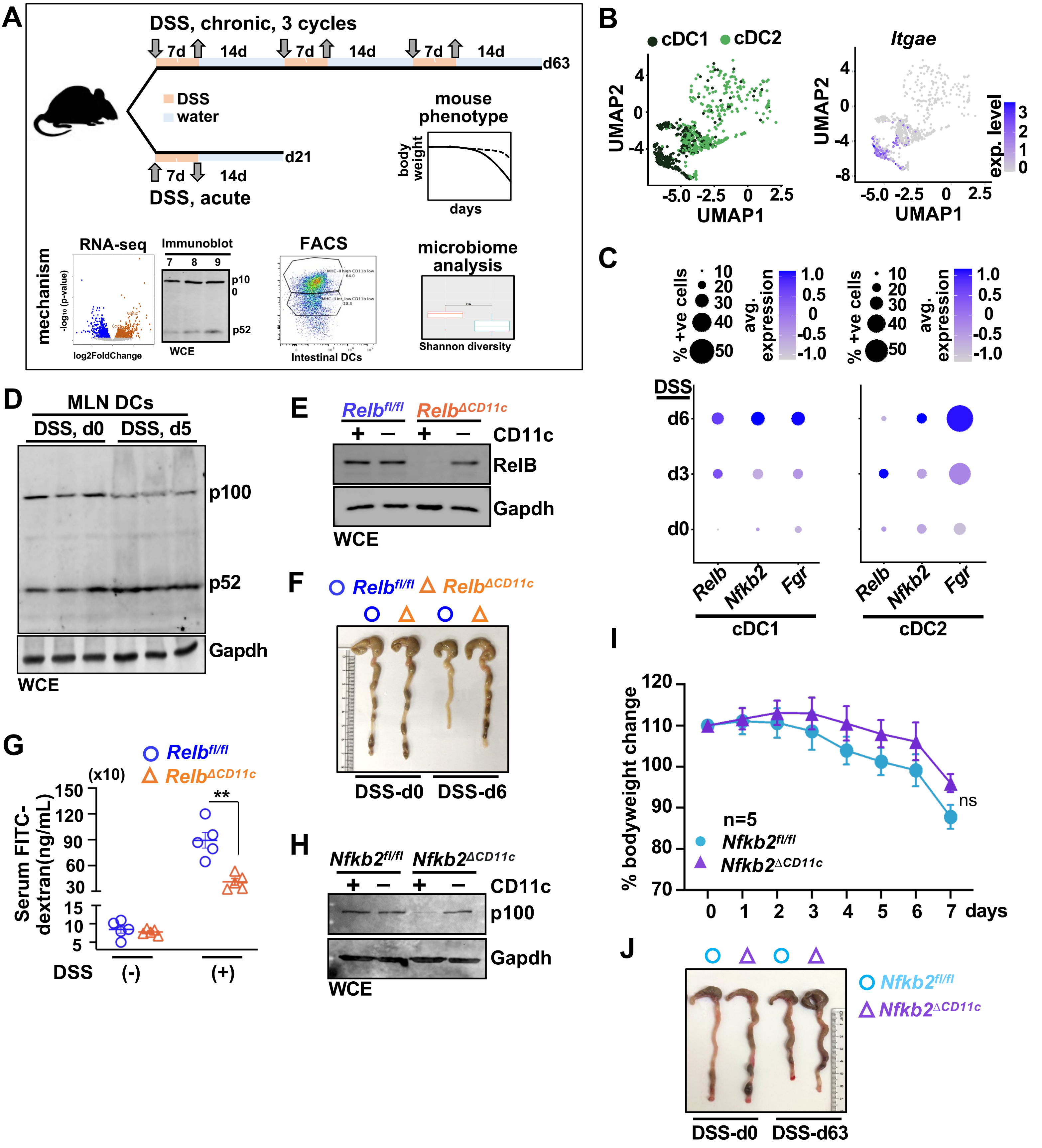


**Figure S1: Studying DSS-induced experimental colitis in mice.** **(A)** Experimental schema for DSS-induced acute and chronic colitis in mice. **(B)** Feature plot depicting the DC-subsets (left panel) and CD103 expression (right panel) in the DC cluster indicated in Fig1A. **(C)** Dotplot revealing the expression of indicated genes in cDC1 (left) and cDC2 (right). Publicly available mouse single-cell RNA-seq data was analysed (GSE148794). **(D)** Immunoblot showing abundance of p52/p100 in MLN-DCs sorted from untreated mice or those administered with 1.5% DSS in drinking water for 5 days. **(E)**Immunoblots revealing the abundance of RelB in CD11c$^{+}$ or CD11c$^{-}$ splenocytes from *Relb$^{fl/fl}$* and *Relb$^{\Delta CD11c}$* mice. **(F)** Representative colon images from *Relb$^{fl/fl}$* and *Relb$^{\Delta CD11c}$* mice subjected to 1.5% acute DSS treatment. **(G)** Dot plot revealing the serum concentration of FITC-dextran in *Relb$^{fl/fl}$* and *Relb$^{\Delta CD11c}$* mice left untreated or subjected to acute DSS treatment; FITC-dextran was gavaged orally 6h prior to serum collection **(H)** Immunoblot showing p100/*Nfkb2* in CD11c$^{+}$ or CD11c$^{-}$ splenocytes from *Nfkb2$^{fl/fl}$* and *Nfkb2$^{\Delta CD11c}$* mice. **(I)** In the acute colitis regime, *Nfkb2$^{fl/fl}$* and *Nfkb2$^{\Delta CD11c}$* mice were treated with 1.5% DSS and evaluated for body weight changes for 7 days **(J)** Representative colon images from *Nfkb2$^{fl/fl}$* and *Nfkb2$^{\Delta CD11c}$* mice subjected to chronic-DSS treatment. Data represent mean ± SEM. For statistical analysis, two-tailed Student's t-test was performed. ns, not significant.

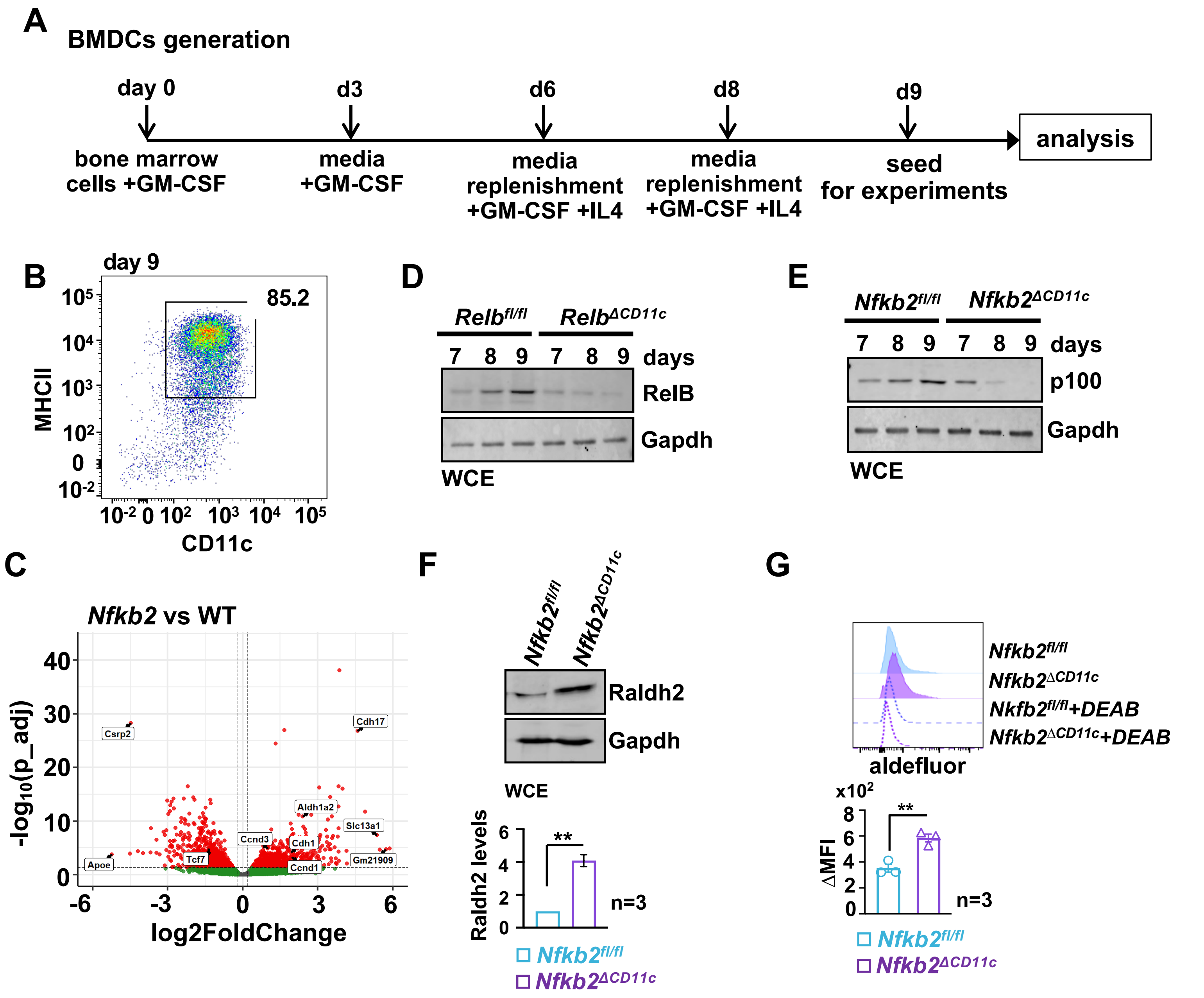


**Figure S2: Studying WT and non-canonical NF-κB-deficient BMDCs. (A)** Schema depicting generation BMDCs *ex vivo*. **(B)** Representative FACS plot showing the percentage of $CD11c^{+}MHCII^{hi/int}$ cells generated on day 9 in BMDC-differentiating culture *ex vivo*. Bone marrow cells from WT C57/BL6 mice were used. **(C)** Volcano plot comparing WT and *Nfkb2*$^{-/-}$ BMDCs for the global gene expression. **(D)** and **(E)** Immunoblot analyses comparing BMDCs derived from *Relb*$^{fl/fl}$ and *Relb*$^{\Delta CD11c}$ mice **(D)** or *Nfkb2*$^{fl/fl}$ and *Nfkb2*$^{\Delta CD11c}$ **(E)** mice for the expression of RelB **(D)** or p100 **(E)**, respectively. **(F)** Representative immunoblot revealing the abundance of Raldh2 in *Nfkb2*$^{fl/fl}$ and *Nkfb2*$^{\Delta CD11c}$ BMDCs. Barplot below represent quantified band intensities. **(G)** Histograms (top) showing Raldh enzymatic activity measured by Aldefluor assay in *Nfkb2*$^{fl/fl}$ and *Nkfb2*$^{\Delta CD11c}$ BMDCs. Data from independent experiments are presented in a barplot below.

Data represent mean ± SEM. For statistical analysis, two-tailed Student's t-test was performed. *P < 0.05; **P < 0.01; ***P < 0.001; ns, not significant.

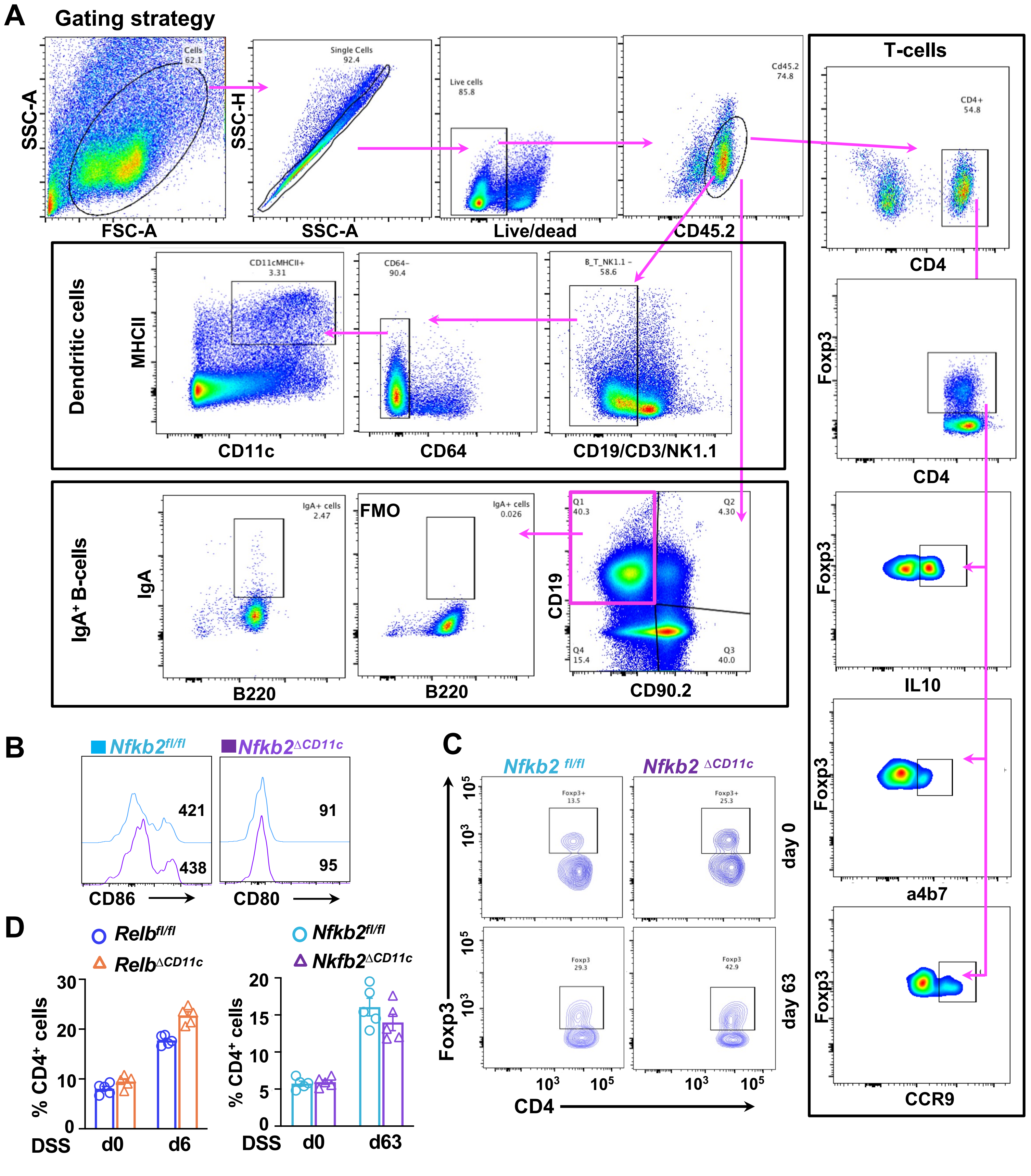



Figure S3: Investigating colonic lamina propria immune cells in mice. (A) Representative FACS plots showing the gating strategy for analyzing the frequencies of dendritic cells, T cell subsets and B-cells in the lamina propria. Fluorescence minus one(FMO) was used as a gating control for gating IgA+ cells. (B) Histograms comparing CD80 and CD86 levels in MLN or LP DCs from Nfkb2$^{fl/fl}$ and Nfkb2$^{\Delta CD11c}$ mice. Corresponding MFI values are indicated in the graph. (C) Flow cytometry analyses showing the frequency of FoxP3+ Tregs in LP or MLNs of Nfkb2$^{fl/fl}$ and Nfkb2$^{\Delta CD11c}$ mice, untreated or subjected to chronic DSS treatment. (D) Barplot revealing the frequency of CD4$^{+}$ T cells as a percentage of CD45.2$^{+}$ cells in Relb$^{fl/fl}$ and Relb$^{\Delta CD11c}$ (left) or Nfkb2$^{fl/fl}$ and Nfkb2$^{\Delta CD11c}$ (right) mice (n=5). Data represent mean $\pm$ SEM. For statistical analysis, two-tailed Student's t-test was performed. *P < 0.05; **P < 0.01; ***P < 0.001; ns, not significant.

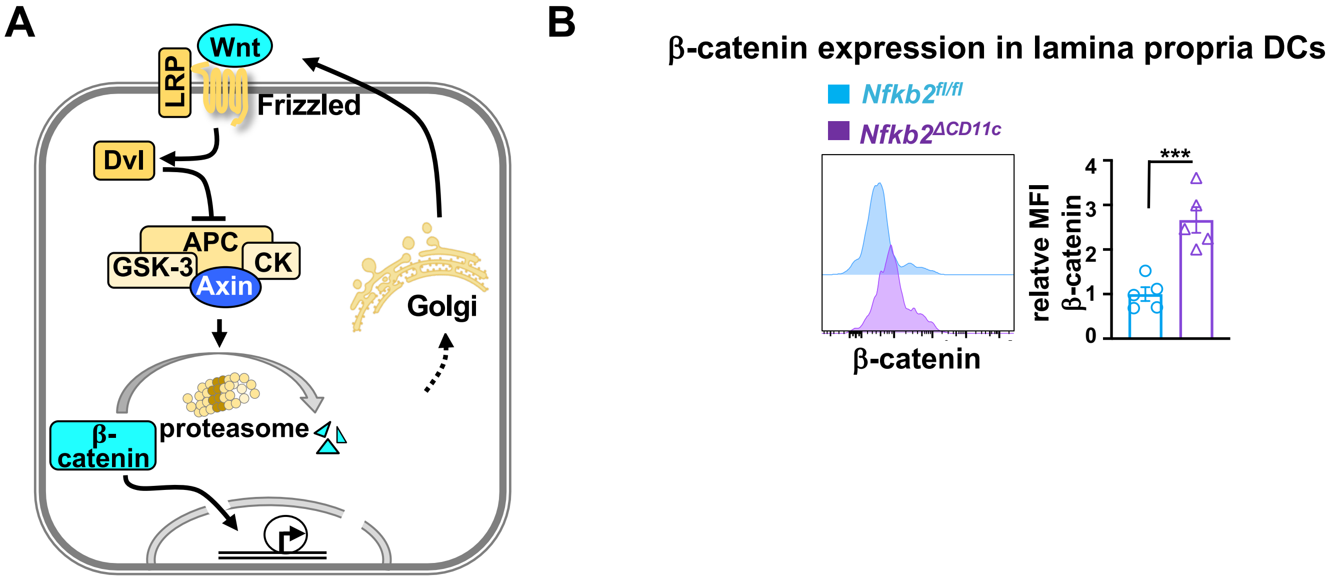


**Figure S4: Investigating crosstalk between β-catenin and non-canonical NF-κB signaling in DCs. (A)** A cartoon describing β-catenin-regulatory mechanisms. **(B)** Representative histogram comparing MLN DCs from *Nfkb2*$^{fl/fl}$ and *Nfkb2*$^{\Delta CD11c}$ mice for the expression of β-catenin.Corresponding quantified data showing relative MFI for β-catenin expression has also been presented in a barplot (right). Data represent mean $\pm$ SEM. For statistical analysis, two-tailed Student's t-test was performed. *P < 0.05; **P < 0.01; ***P < 0.001

### Supplementary Table S1- Prototypic gene signatures for cell types/subsets

| Cell type/subset | Signature genes |
|---|---|
| Dendritic cells (adopted from Xu et al., 2019[9]) | CD74, H2AB1, H2EB1, H2AA, TMSB4X, CST3, CD52, H2DMA, PSAP, CRIP1, SH3BGRL3, TYROBP, LSP1, FCER1G, CORO1A, MPEG1, IRF8, PLBD1, GPX1, LGALS3, SRGN, CTSS, ALOX5AP, IFI30, GM2A, XCR1, CLEC9A, ITGAX, CCR2, CLEC7A, CD83, CD86, BATF3, CCR7, ZBTB46, BCL11A, FLT3, DPP4, CD8A, CD14, CSF1R, CX3CR1, FCGR3, FCGR1, CD207, CD209A, ITGAM, SIRPA, CD4, IRF4, TBX21, MGL2, ESAM, DTX1, RBPJ, SIGLECH, IRF7, TCF4, BST2, IL23A, CXCR3, CCR6 |
| Macrophages (adopted from Xu et al., 2019[9]) | CCL8, APOE, C1QB, C1QC, LYZ2, C1QA, CTSB, SELENOP, ITM2B, FTH1, H2-D1, TMSB4X, TYROBP, CTSS, PF4, B2M, FTL1, LGMN, GRN, FCGR3, CSF1R, CST3, LAPTM5, MRC1, WFDC17, F13A1, SERINC3, FCER1G, CTSC, CD74, CLEC4F, CD5L, VSIG4, FCNA, PSAP, MAFB, CD68, CD163, CD163L1, MERTK, CD209, STAB1, SLCO2B1,MMP12, MMP14, F480, ITGAM, CX3CR1, FCGR1, ADGRE1, CD64 |
| Type-I Classical DC (cDC1) | XCR1, BATF3, IRF8, CD8A, ITGAE, FLT3, CLEC9A, LY75 |
| Type-II Classical DC (cDC2) | ITGAM, SIRPA, IRF4, CSF1R, ZBTB46, MARCO, CLEC4 |
