## Supplementary material for "Non-canonical NF-κB signaling promotes intestinal inflammation by restraining the tolerogenic β-catenin-Raldh2 axis in dendritic cells": Key resource table.pdf

**Table1- Reagents**

| Reagent | Source | Identifier |
|---|---|---|
| **Antibodies** | | |
| Anti-RelA rabbit polyclonal | Santa Cruz Biotechnology | sc-372; AB_632037 |
| Anti-RelB rabbit polyclonal | Cell Signaling Technology | 4922S; AB_2179173 |
| Anti-mouse p52/p100 rabbit polyclonal | Generated in the Systems Immunology Lab | Mouse p52/p100 sequence used: DNCY DPGLDGIPEYDD |
| Anti-Gapdh rabbit polyclonal | Cell Signaling Technology | 2118S; AB_10698756 |
| Anti-active β-catenin rabbit monoclonal | Cell Signaling Technology | 8814S; AB_11127203 |
| Anti-Raldh2 rabbit monoclonal | Cell Signaling Technology | 83805S; AB_2800032 |
| Anti-phospho-GSK-3β rabbit polyclonal | Cell Signaling Technology | 9336S; AB_331405 |
| Anti-GSK-3β rabbit polyclonal | Cell Signaling Technology | 9315S; AB_490890 |
| Anti-Axin1 rabbit monoclonal | Cell Signaling Technology | 2087S; AB_2274550 |
| Anti-Notch2 rabbit monoclonal | Cell Signaling Technology | 5732T;AB_10693319 |
| Anti-β-catenin Rabbit polyclonal | Invitrogen | 71-2700; AB_2533982 |
| Anti-IRF4 rabbit polyclonal | Thermo Scientific | PA5-21144; AB_11152871 |
| Goat anti-Rabbit IgG-Cy5 secondary | Cytiva | PA45011; AB_772205 |
| Rabbit Trueblot Anti-Rabbit IgG HRP | Rockland | 18-8816-33; AB_2610848 |
| Anti-CD45.2 (104) BUV563 | eBiosciences | 365045482; AB_2925379 |
| Anti-CD25 (PC61.5) BUV661 | eBiosciences | 376025182; AB_2925446 |
| Anti-IL17A (17B7) BUV805 | eBiosciences | 368717782; AB_2896169 |
| Anti-CD197(CCR7) 4B12 eF450 | eBiosciences | 48197182; AB_1944351 |
| Anti-CD4(GK1.5) BUV737 | eBiosciences | 367004182; AB_2895921 |
| Anti-integrin a4b7 PE (DATK32) | eBiosciences | 12588782; AB_657803 |
| Anti-MHC II (M5/114.15.2) PE-Cy7 | eBiosciences | 25532182; AB_10870792 |
| Anti-CD64 (X54-5/7.1) PerCP-e710 | eBiosciences | 46064182; AB_2735016 |
| Anti-CD19 (1D3) BV780 | eBiosciences | 780109382; AB_2925722 |
| Anti-CD3E (145-2C11) BV780 | eBiosciences | 78003182; AB_2784894 |
| Anti-NK1.1(PK136) BV780 | eBiosciences | 78594182; AB_2744923 |
| Anti-Foxp3(150D/E4) FITC | eBiosciences | 11577382; AB_465243 |
| Anti-F4/80(BM8) FITC | eBiosciences | 123107;AB_103762287 |

| | | |
|---|---|---|
| Anti-CD11b (M1/70) BV 510 | BioLegend | 101245; AB_2561390 |
| Anti-CD11c (N418) PerCP | BioLegend | 117236; AB_925727 |
| Anti-β-catenin1(15B8) PE | BioLegend | 862604;AB_2832863 |
| Anti-CCR9 (CW1.2) PE-Cy7 | BioLegend | 128712; AB_10901176 |
| Anti-IFNγ (XMG1.2) APC-Cy7 | BioLegend | 505850; AB_2616698 |
| Anti-B220 (RA3-6B2) FITC | BD Pharmingen | 553088; AB_394618 |
| Anti-IgA (mA-6E1) PE | eBiosciences | 12-4204-82;AB_465917 |
| **Chemicals and Peptides** | | |
| DSS (36,000-50,000 MW) | MP Biomedicals | 160110 |
| FITC-Dextran (MW 3000 - 5000) | Sigma-Aldrich | 60842-46-8 |
| Collagenase, Type 4 | Gibco | 17104019 |
| DNase I | Worthington | LS002139 |
| LPS from E. coli serotype O55:B5 | Enzo Life Sciences | ALX-581-013-L002 |
| Porcn Inhibitor-II, C59-inhibitor | Merck | SKU5004960001 |
| Wnt-β-catenin signaling inhibitor -Fzm1 | Merck | 534358 |
| BMS493 (RAR inhibitor) | MedChem Express | HY-108529 |
| β-catenin/Tcf InhibitorIII, iCRT3 | Sigma-Aldrich | 219332 |
| Recombinant mouse TGFβ1 | BioLegend | 763104 |
| anti-CD3 | Thermo Scientific | 16-0032-86;AB_467057 |
| anti-CD28 | Thermo Scientific | 16-0281-86;AB_468923 |
| recombinant mouse GM-CSF | Miltenyi Biotec | 130095739 |
| recombinant mouse IL-4 | Miltenyi Biotec | 130097757 |
| **Commercial Kits** | | |
| RNeasy Mini Kit (250) | Qiagen | 74106 |
| Primescript 1$^{st}$ strand cDNA synthesis kit | Takara Bio | 6110B |
| Foxp3/Transcription Factor Staining Buffer | Invitrogen | 00-5523-00 |
| ALDEFLUOR Kit | Stem cell technologies | 01700 |
| Leukocyte activation kit | BD Biosciences | 550583 |
| Naive CD4-T cell isolation kit | Miltenyi Biotec | 130 104 453 |
| Live/dead fixation yellow dye | Invitrogen | L34968 |
| QIAamp PowerFecal Pro DNA kit | Qiagen | 51840 |
| Mouse IgA Uncoated ELISA kit | Invitrogen | 88-50450-22 |
| **Mice models** | | |
| *Nfkb2*$^{-/-}$ | small animal facility, NII | N/A |
| *Relb*$^{fl/fl}$ mice | Jackson Laboratories | 028719 |
| *Nfkb2*$^{fl/fl}$ mice | Jackson Laboratories | 028720 |
| *Ctnnb1* $^{fl/fl}$ mice (floxed for β-catenin gene) | generous gift from Prof. Amitabha Bandopadhyay, IIT Kanpur | N/A |
| *Itgax*-Cre mice (CD11c-Cre) | Jackson Laboratories | 008068 |

### Table2-Primers for qRT-PCR (mouse genes)

| Gene | Forward (5' – 3') | Reverse (3' – 5') |
|---|---|---|
| *Il1b* | AACCTGCTGGTGTGTGACGTTC | CAGCACGAGGCTTTTTGTTGT |

| | | |
|---|---|---|
| *Cxcl10* | ACCAACCACCAGGCTAGA | GCGTCACACTCAAGCTCT |
| *Aldh1a2* | ATGGGTGAGTTTGGCTTACG | GGTTCATTGGAAGGCAGAAA |
| *Il10a* | CTAACCGACTCCTTAATGC | AATCACTCTTCACCTGCTC |
| *Actin* | CCAACCGTGAAAAGATGAC | GTACGACCAGAGGCATACAG |
| *Ctnnb1* | GTTCGCCTTCATTATGGACTGCC | ATAGCACCCTGTTCCCGCAAAG |
| *Axin1* | CACCCAGAAGCTGCTATTGGAGA | CCAGGGCATAGCCAGAGTTGA |
| **Bacterial taxa primers** | | |
| *Actinobacteria* | CGCGGCCTATCAGCTTGTTG | ATTACCGCGGCTGCTGG |
| *Firmicutes* | GGAGYATGTGGTTTAATTCGAAGCA | AGCTGACGACAACCATGCAC |
| *Bifidobacterium* | TCGCGTCCGGTGTGAAAG | CCACATCCAGCATCCAC |
| *Enterobacter* | GTGCCAGCMGCCGCGGTAA | GCCTCAAGGGCACAACCTCCAAG |
| *SFB* | GACGCTGAGGCATGAGAGCA | GACGGCACGGATTGTTATTC |
| *Sutterella* | CGCGAAAAACCTTACCTAGCC | GACGTGTGAGGCCCTAGCC |
| *Prevotella* | CACGGTAAACGATGGATGCC | GGTCGGGTTGCAGACC |
| *β-Proteobacteria* | TCACTGCTACACGYG | ACTCCTACGGGAGGCAGCAG |
| *Universal 16s* | TCCTACGGGAGGCAGCAGT | GGACTACCAGGGTATCTAATCCTGTT |

**Table3-Software/computational tools**

| Software | Source |
|---|---|
| DESeq2 package | https://bioconductor.org/packages/release/bioc/html/DESeq2.html [53] |
| fgsea package | https://bioconductor.org/packages/release/bioc/html/fgsea.html [54] |
| Integrative Genomics Viewer v2.16.2 | https://igv.org/app/ [55] |
| MEME Suite | https://meme-suite.org/meme/tools/meme [56] |
| AUCell package | https://bioconductor.org/packages/release/bioc/html/AUCell.html [57] |
| Seurat package | https://satijalab.org/seurat/reference/seurat-package [58] |
| fastQC | https://www.bioinformatics.babraham.ac.uk/projects/fastqc/ |
| Paired-End reAd mergeR (PEAR) | https://www.h-its.org/downloads/pear-academic/ [59] |
| QIIME2-2022.2 | https://qiime2.org/ [60] |
| SILVA SSU 138 | https://www.arb-silva.de/documentation/release-1381/ [61] |
| ampvis R-package | https://kasperskytte.github.io/ampvis2/articles/ampvis2.html |
| ggplot2 package | https://ggplot2.tidyverse.org/ [62] |
| Biorender | https://www.biorender.com/ |
